# Runx1 promotes scar deposition and inhibits myocardial proliferation and survival during zebrafish heart regeneration

**DOI:** 10.1101/799163

**Authors:** Jana Koth, Xiaonan Wang, Abigail C. Killen, William T. Stockdale, Helen G. Potts, Andrew Jefferson, Florian Bonkhofer, Paul R. Riley, Roger Patient, Berthold Göttgens, Mathilda T.M. Mommersteeg

**Affiliations:** Department of Physiology, Anatomy and Genetics, University of Oxford, South Parks Road, Oxford OX1 3PT, United Kingdom; MRC Molecular Haematology Unit, Weatherall Institute of Molecular Medicine, John Radcliffe Hospital, University of Oxford, United Kingdom; Department of Haematology, Wellcome and MRC Cambridge Stem Cell Institute, University of Cambridge, Cambridge, United Kingdom; Micron Advanced Bioimaging Unit, Department of Biochemistry, South Parks Road, Oxford OX1 3QU, United Kingdom

**Keywords:** zebrafish heart regeneration, Runx1, fibrinolysis, smooth muscle

## Abstract

Runx1 is a transcription factor that plays a key role in determining the proliferative and differential state of multiple cell-types, during both development and adulthood. Here, we report how *runx1* is specifically upregulated at the injury site during zebrafish heart regeneration, but unexpectedly, absence of *runx1* results in enhanced regeneration. Using single cell sequencing, we found that the wild-type injury site consists of Runx1-positive endocardial cells and thrombocytes that induce expression of smooth muscle and collagen genes without differentiating into myofibroblasts. Both these populations are absent in *runx1* mutants, resulting in a less collagenous and fibrinous scar. The reduction in fibrin in the mutant is further explained by reduced myofibroblast formation and by upregulation of components of the fibrin degradation pathway, including plasminogen receptor Annexin 2A as well as downregulation of plasminogen activator inhibitor *serpine1* in myocardium and endocardium, resulting in increased levels of Plasminogen. In addition, we find enhanced myocardial proliferation as well as increased myocardial survival in the mutant. Our findings suggest that Runx1 controls the regenerative response of multiple cardiac cell-types and that targeting Runx1 is a novel therapeutic strategy to induce endogenous heart repair.

## Introduction

Heart regeneration potential varies considerably between species as well as with age. While zebrafish, neonatal mouse and neonatal human hearts can replace dead or lost cardiomyocytes rapidly with new heart muscle (1–5), medaka (6, 7), cave fish (8), as well as adult mice and human hearts (9) show only poor repair. Numerous studies are, therefore, looking into the underlying principles and mechanisms that promote, or prevent, effective cardiac regeneration to establish a basis for therapeutic intervention (10). In models of successful regeneration, remaining cardiomyocytes have been shown to proliferate and replace the scar tissue with new heart muscle (2, 11–13). This is dependent on a fine balance of interaction with other cell-types, including the epicardium and endocardium (10, 14). Due to the complexity of this interaction, we still lack a clear understanding of how the scar tissue can be broken down and replaced by proliferating myocardial cells. Here, we report a role for Runx1 in regulating the delicate balance between scar degradation and myocardial regeneration.

Runx transcription factors, which hetero-dimerise with core binding factor β (CBFβ), are transcription factors that can function as activators as well as repressors and, as such, are important regulators of lineage-specific cell fate. Runx1 (also known as acute myeloid leukaemia 1 protein (AML1) or core-binding factor subunit alpha-2 (CBFA2)) is a master transcription factor for determining the proliferative and differential state of multiple cell-types, during both development and adulthood. Runx1 is most studied for its role in endothelial-to-haematopoietic transition during haematopoiesis in development (15–20) and as a well-known fusion oncogene (21, 22). Relatively little is known about the role of Runx1 in skeletal and heart muscle. It has been shown that Runx1 is important in skeletal muscle stem cell (SC) proliferation and its levels can affect the proliferative timing and thus the regenerative capacity of skeletal muscle cells (23). In the heart, Runx1 is expressed in neonatal mouse cardiomyocytes and is upregulated in zebrafish, adult mouse, rat and human cardiomyocytes after injury (24–28). Conditional *Runx1* deficiency in mouse cardiomyocytes has been demonstrated to protect the mouse against the negative consequences of cardiac remodelling after myocardial infarction (29). Although no changes in injury size were found between myocardial conditional *Runx1* knock-out and control mice, the remaining cardiomyocytes displayed improved calcium handling, accompanied by improved wall thickness and contractile function compared to wild-type (29). However, as the knockout was cardiomyocyte specific, the involvement of other cardiac cell-types was not investigated. In contrast to mouse, where constitutive *Runx1* deletion is embryonically lethal, zebrafish *runx1^W84X^* mutants (30) are homozygote viable adults, allowing us to investigate the role of *runx1* loss of function during zebrafish heart repair down to the single cell level.

We show that Runx1 has important roles in the response of various cell-types to injury, including thrombocytes, the epicardium, endocardium and myocardium. Thrombocytes are the fish equivalent of platelets and important for blood clotting, with the difference that these are nucleated cells (31). While removal of *runx1* leads to unique cell-type specific responses, in combination, these tip the balance towards faster scar degradation and enhanced myocardial regeneration. Runx1 not only controls myocardial survival and proliferation, it also regulates the composition and degradation of extracellular matrix at the wound site. Thrombocytes and endocardial cells that normally express smooth muscle and collagen genes are missing in the mutant, not only changing the cellular composition of the wound, but also strongly reducing the amount of collagen and fibrin deposition. The epicardium shows reduction in the level of smooth muscle and collagen genes in the *runx1* mutant, on top of which there is a strong reduction in the number of myofibroblasts formed. Degradation of the wound is further enhanced by increased fibrinolysis, allowing invasion of highly proliferative cardiomyocytes and improved heart regeneration. Collectively this leads to enhanced heart regeneration in the absence of *runx1* and identifies Runx1 inhibition as a potential therapeutic target to improve cardiac repair in mammals.

## Results

### Runx1 becomes widely expressed in zebrafish hearts after injury

To evaluate *runx1* expression in the adult heart we induced cryo-injury using a liquid nitrogen cooled probe in the *Tg(BAC-Runx1P2:Citrine)* zebrafish line, in which cytoplasmic Citrine fluorescence is placed under the control of the *runx1* P2 promotor (32). The P2 promoter is the main one of 2 *runx1* promoter regions known to drive expression in definitive hematopoietic stem cells (HSCs) in the dorsal aorta during development (20), however, its expression in the adult heart is unknown. In the uninjured heart, Runx1-Citrine expression was sparse but present in a small number of cells spread throughout the heart, mostly blood cells (Fig. 1a-a’). However, after injury, expression became much more widespread: one day post cryo-injury (dpci), a large collection of bright Citrine positive cells was present in the injury site (Fig. 1b-b’), indicating the presence of Citrine-positive blood cells in the wound. In addition to the blood cells, other cell populations started to express Citrine, including cells within the epicardium all around the heart (arrowheads, Fig. 1b). Additionally, weak expression of Citrine was observed in cardiomyocytes bordering the injury site (Fig. 1b’-b”, insert). Three days after injury, Citrine expression in these cell-types was even more pronounced, especially within the endocardium specifically near the injury site (arrowheads, Fig. 1c-c”). Moreover, at this time-point, myocardial cells surrounding the injury site strongly expressed Citrine as shown by overlapping expression of Citrine with the myocardial marker MF20 (Fig. 1c-c”, insert). This pattern was maintained at 7dpci, but started to taper-off around 14dpci (Supp Fig. 1a-a”, Fig. 1c-d”). Even in sham operated hearts, in which the ventricle was only exposed to a room-temperature probe, Citrine expression was upregulated in both the epicardium (arrowheads) and myocardium, but not endocardium (Supp Fig. 1b-b”). To verify the cell-type specific expression of Citrine, we confirmed overlapping expression with different cell-type specific markers. The bright blood cell population present in the wound at 1dpci was also highly positive for *itga2b*, which is a marker for nucleated thrombocytes (arrowheads, Fig. 1e-e’) (33). Additionally, we found Citrine overlapping with leukocyte marker LyC, endothelial/endocardial marker ERG1 and epicardial/fibroblast marker *tcf21* (Supp Fig. 1c-e) at 3dpci. As *runx1* expression was analysed by visualisation of a transgene, we also checked if transgene expression followed the same pattern as endogenous runx1 RNA using RNA-scope *in situ* hybridisation (34). Runx1 RNA and Citrine expression showed clear overlapping expression patterns, with RNA present in the Citrine positive epicardium, myocardium and endocardium (Supp Fig. 2a-c) after injury. Runx1 RNA was not or very low expressed in the Citrine-negative myocardium of the rest of the ventricle (Supp Fig. 2b-c, asterisks). To summarise, *runx1* expression becomes strongly upregulated in several cell populations of the heart after injury, in the myocardium, endocardium and epicardium surrounding the wound area. This upregulation of runx1-Citrine after cryo-injury suggests a role for *runx1* in multiple cell-types during heart regeneration.

**Fig. 1.**
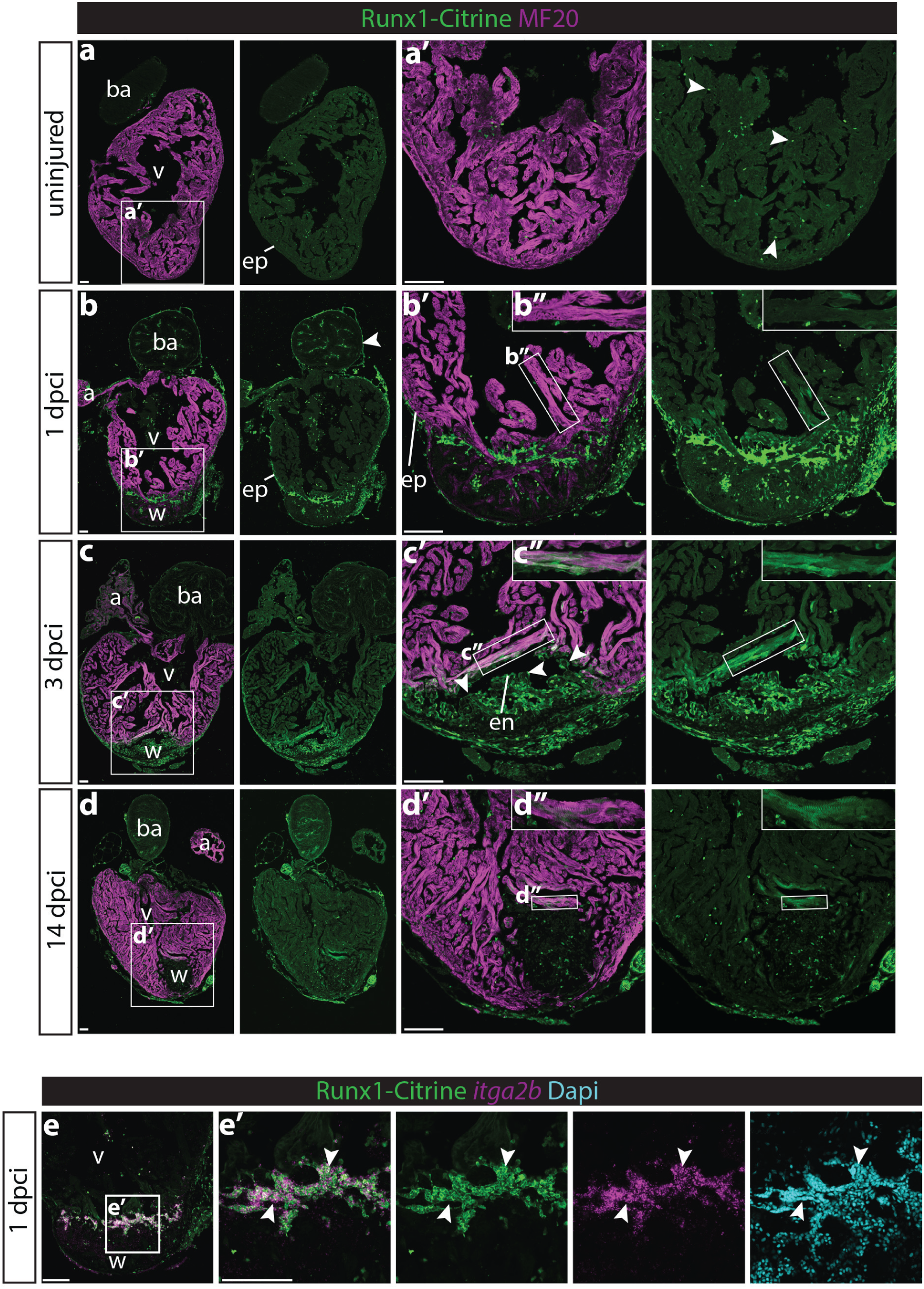
Runx1-Citrine becomes strongly expressed in the heart after cryo-injury. a-d”, immunohistochemistry for Runx1-Citrine (GFP antibody) and myocardial marker MF20 at different time points after cryo-injury. a-a’, Citrine expression in the uninjured hearts was confined to a small number of cells scattered around the heart (arrowheads). b-b”, at 1 dpci the epicardium was Citrine-positive (arrowheads), as well as bright blood cells within the wound and dim expression of Citrine overlapping with MF20 (b”). c-c”, at 3 dpci, the epicardium, endocardium (arrowheads) and other wound cells were positive for Citrine. Also the myocardium in the border zone next to the wound was highly Citrine-positive (c”). d-d”, expression of Citrine diminishes at 14 dpci. But expression is still visible, especially in the myocardium (d”). e-e’, *in situ* hybridisation for *itga2b* with immunohistochemistry for Runx1-Citrine and nuclear marker Dapi. Arrowheads point to overlap of Runx1-Citrine with itga2b mRNA indicating that thrombocytes are positive for Runx1-Citrine. a, atrium; ba, bulbus arteriosus; dpci, days post cryo-injury; en, endocardium; ep, epicardium; v, ventricle; w, wound. Scale bars depict 100 μm.

**Fig. 2.**
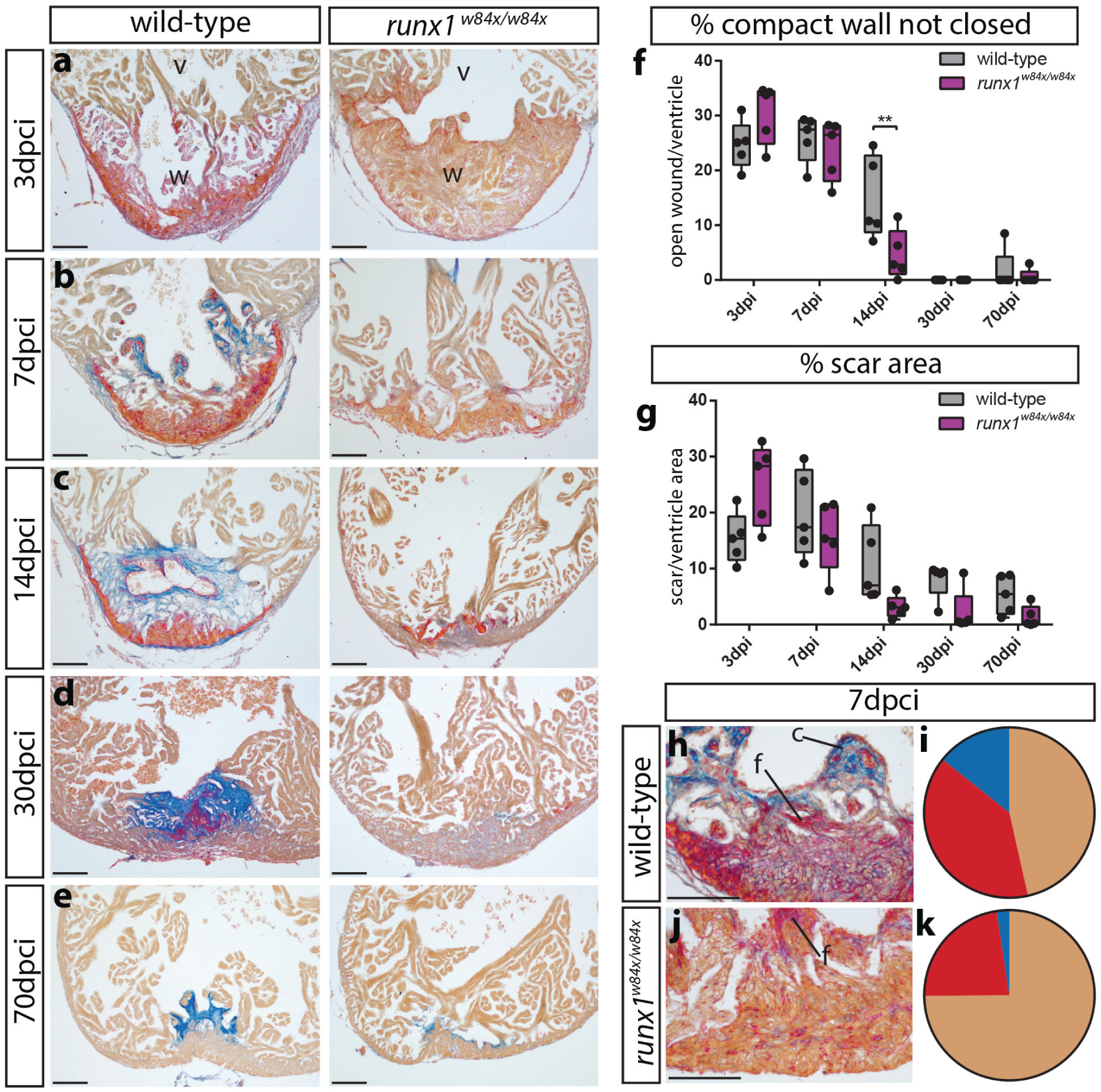
Different wound composition and faster regeneration in *runx1* mutant compared to wild-type hearts. a-e, AFOG staining of wild-type and *runx1* mutant ventricles at 5 different time points after injury. f-g, quantification of the difference in wound size between the wild-type and mutant at the different time points, measured by percentage of the compact wall not yet closed (f) and the % of scar area compared to total ventricle area (g). n=5 per time point, two-way ANOVA with Sidak test. h-i, quantification of differences in wound composition between the fish at 7dpci, n=5. c, collagen; f, fibrin; v, ventricle; w, wound. Scale bars depict 100 μm.

### Enhanced regeneration in *runx1* mutants compared to wild-type zebrafish

Based on the observed expression of Runx1 after injury, we questioned if heart regeneration is affected in absence of *runx1*. We used the global *runx1^W84X/W84X^* mutant, which has a truncation mutation, leading to complete loss of function (35). After performing cryo-injury on *runx1* wild-type and mutant fish we isolated the hearts at 5 different time-points, from 3 to 70 days post injury. Acid Fuchsin Orange G staining (AFOG, labelling collagen in blue, fibrin in bright red and myocardium/blood cells in orange), showed a clear difference between wild-type and mutant hearts (Fig. 2a-k). At 3dpci, the injury site was clearly visible in both fish, but the wild-type hearts showed a much more extensive deposition of red fibrin compared to the mutants (Fig. 2a). As well as less fibrin deposition, we also observed reduced collagen (blue) deposition at 7dpci in the mutants (Fig. 2b, h-k). While the wild-type wound consisted on average of 39.2% fibrin (red) blood clot and 14.3% collagen (blue), *runx1* mutant hearts had around 22.7% fibrin (red) and 2.4% collagen (blue) labelling. Despite these differences in wound composition, comparison of the wound size did not show any significant differences between the wild-type and mutant hearts at 3 and 7dpci (Fig. 2a-b, f-g). However, at 14dpci, there was a significantly stronger decrease in open wound length in the mutant compared to the controls, indicating a faster resolution of the lesion (Fig. 2c, f-g). At 30dpci both mutants and controls had closed the compact myocardial wall over the wound, but the remaining scar was less visible in the mutants, with blue collagen mainly present in between the regenerated trabeculae (Fig. 2d). The difference was still visible at 70dpci, with trace amounts of blue collagen label present in the mutants. These data show that the *runx1* mutants have a significantly larger area of their compact wall closed at 14dpci compared to wild-types and deposit a different extracellular matrix after heart injury compared to wild-types.

### Significant increase in Runx1-Citrine positive endocardial cells after injury

Since we observed strong expression of Runx1-Citrine in the endocardium, which plays crucial roles during regeneration and scar formation (10, 36, 37), we next analysed this in more detail (Fig. 3a-f). We crossed the *Tg(BAC-Runx1P2:Citrine)* with the *Tg(kdrl:Hsa*.*HRAS-mCherry)* line, which in combination label Runx1-Citrine endothelial/endocardial cells with membrane mCherry fluorescence (38), and initially observed very few Citrine-mCherry double positive cells in intact or sham-operated hearts (Fig. 3e). At 1dpci, we observed a significant increase in mCherry positive cells in the wound area with a flat endocardial cell morphology and dim Citrine expression compared to the bright Citrine-positive blood cells (Fig. 3a’, e). Double positive cells were most clearly visible at 3dpci (Fig. 3b-b’, e), with a rounder cell morphology, while throughout the remaining intact ventricle away from the wound, only few cells were observed at all stages analysed (ventricle, Fig. 3c-c’, e). This Citrine/mCherry double positive population was still highly present at 7dpci, but decreased towards baseline levels at 14dpci (Fig. 3d-e). The known functions of Runx1 (15–20), combined with its extensive expression pattern in the endocardium, suggests a role for *runx1* and the endocardium in the deposition of the wound extracellular matrix after injury.

**Fig. 3.**
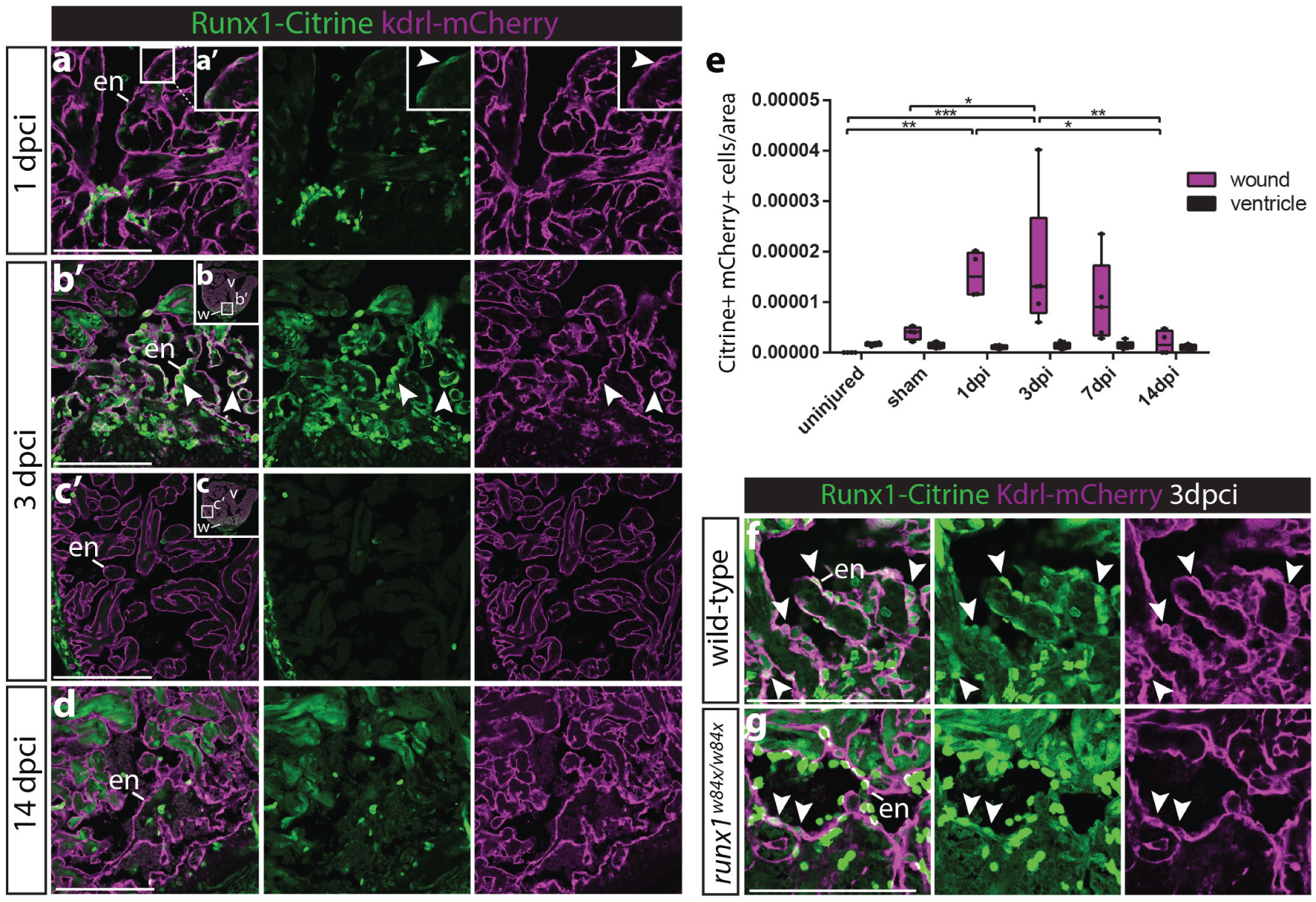
Runx1-Citrine positive endocardial cells appear in the wound after injury. a-g, immunohistochemistry analysis for Citrine and mCherry positive cells in the *Tg(BAC-Runx1P2:Citrine;kdrl:Hsa*.*HRAS-mCherry)* line. a-a’, shows the wound at 1dpci, with the box highlighting the flat and weakly Citrine positive mCherry-positive endocardial cells in the wound. b-c, show well visible and round Citrine-mCherry positive cells in the wound at 3dpci (b-b’), but not further away from the wound (c-c’). d-d’, at 14dpci not many double positive cells are visible anymore. e, quantification of the number of Citrine-mCherry double positive cells in and away from the wound, n4, two-way ANOVA with Tukey’s test. f-g, runx1 mutant wounds have a reduced number of double positive cells. en, endocardium; v, ventricle; w, wound. Scale bars depict 100 μm.

### Single cell sequencing identifies subpopulations of *runx1* expressing cells after injury

To further investigate the function of Runx1 in all *runx1* expressing cell types after injury, with a focus on the endocardium/endothelium, we performed single cell sequencing using the 10x Genomics platform. The *runx1^W84X/W84X^* mutant line was crossed into the *Tg(BAC-Runx1P2:Citrine;kdrl:Hsa*.*HRAS-mCherry)* background and confirmed for preserved BAC-Runx1P2:Citrine expression (Supp Fig. 3a). Citrine expression was overall similar to that in the wild-type heart, including expression in endocardial cells. However, we observed a reduction in the number of Citrine and mCherry double positive cells in the mutant wound (Fig. 3f-g). The ventricles of *runx1* wild-type uninjured, *runx1* wild-type 3dpci and *runx1* mutant 3dpci *Tg(BAC-Runx1P2:Citrine;kdrl:Hsa*.*HRAS-mCherry)* fish were dissociated and FACS sorted (Fig. 4a). FACS sorting for Citrine and mCherry, we found a 4.5-fold increase in double positive cells in wild-type 3dpci hearts compared to wild-type uninjured hearts, confirming our image-based cell counts (Supp Fig. 3b, Fig. 3e). Mutant hearts had only a third of the amount of double positive cells after injury compared to wild-types (Supp Fig. 3b). Before sequencing, we excluded negative cells and combined all single positive and double positive cells, while enriching for the double positive population which might otherwise have been missed during single cell analysis, due to their low numbers. Sequencing and subsequent clustering of all cells combined led to the identification of 27 cell clusters (C) that comprised all the expected cell populations, including endocardial/endothelial cells, myocardial cells, epicardial cells, myofibroblasts, thrombocytes and different leukocyte populations (Fig. 4b-c, supp Fig. 4a). Kdrl/mcherry mRNA positive cells were mainly present in a large group of closely related cell clusters (C0-6), whereas runx1/citrine mRNA-positive cells were as expected present in all clusters, and double positive cells largely grouped in the main *kdrl/mcherry* group (Fig. 4d). In the uninjured heart, endocardial/endothelial cells grouped into 3 main different clusters (C1,3,4), indicating a degree of heterogeneity within these cell populations (Fig. 4e). After injury in both wild-type and *runx1* mutant hearts, 2 large additional endocardial/endothelial cell populations appeared (C0,2) while C3 was reduced in number (arrowheads, Fig. 4e). These injury-specific endocardial populations were highly positive for *serpine1* expression (Supp Fig. 4b), indicating that this population is largely similar to the previously identified highly mobile serpine1-positive endocardial population (36), which was confirmed on sections (Supp Fig. 4c-c’, arrowheads). Although *citrine*-positive populations of neutrophils and macrophages were present in both the mutant and wild-type after injury (C13,15), we observed differences in other blood cell populations and most notably mutant hearts lacked an obvious population of mature thrombocytes and monocytes (C 24-25 and C16, Fig. 4b-e). In contrast, other blood cell clusters unique to the mutant were present (C19, 22, 23), and characterised by highly expressed genes such as *gata2b* or *myb* (Supp Fig. 4d). Analysis of wild-type and mutant tissue sections confirmed the unique presence of these abnormal blood cell populations in the mutant, resulting in an altered leukocyte profile in the wound after injury (Supp. Fig. 4e-h’). These results show on a single cell basis how Runx1 becomes activated after injury with specific cell composition differences between the mutant and wild type hearts.

**Fig. 4.**
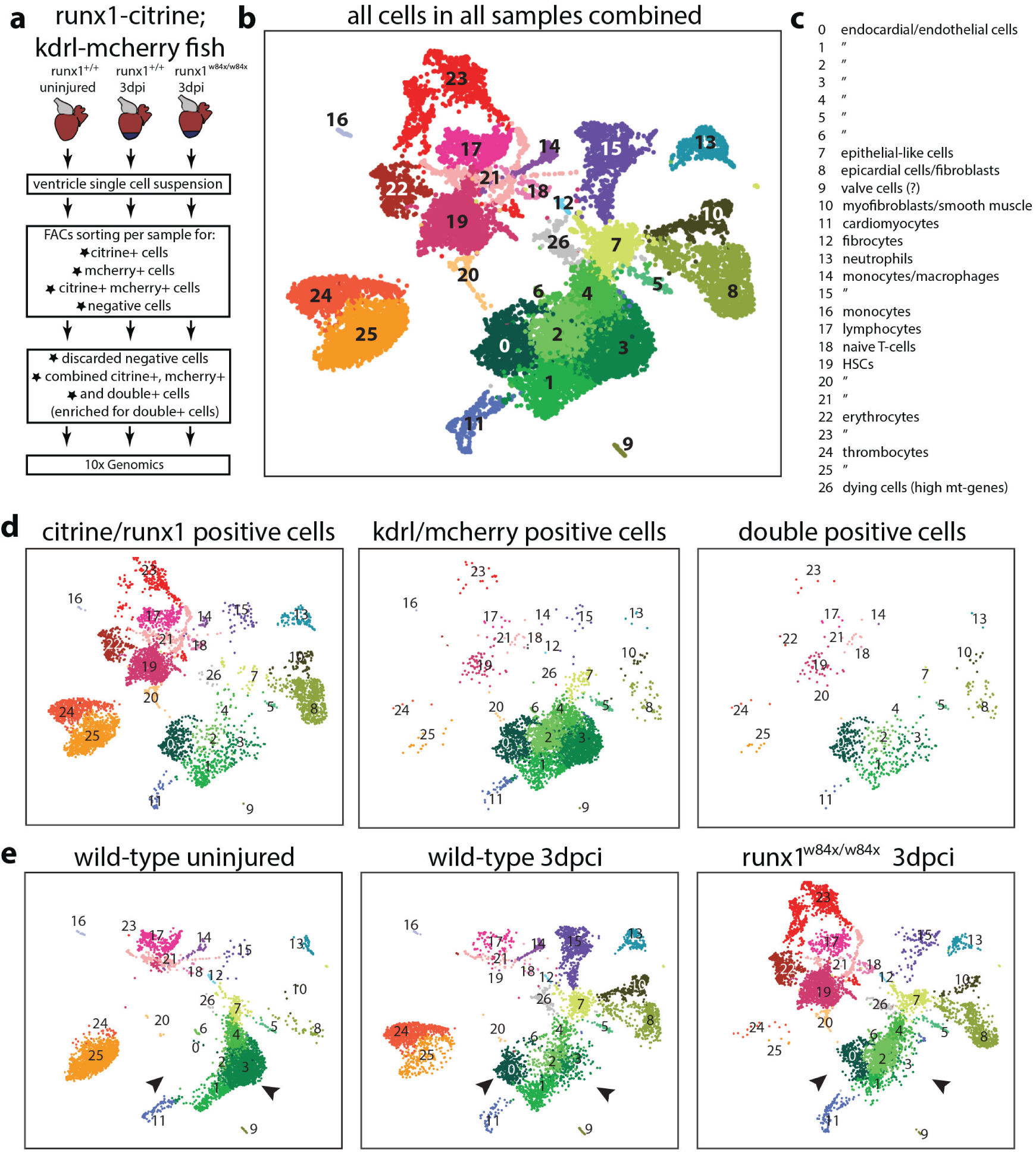
Single cell sequencing of Citrine and mCherry positive cells. a, experimental design of selection of cells for single cell sequencing using the 10x Genomics platform. b, UMAP plot of all cells combined, clustering into 27 different clusters. c, annotation of the different cell clusters. d, UMAP plot separated into *citrine/runx1*-positive cells, *mcherry/kdrl* -positive cells and double positive cells. e, UMAP plot separated into wild-type uninjured cells, wild-type 3dpci cells and *runx1* mutant 3dpci cells. Arrowheads point to the shift in endocardial/endothelial cells, with C0 and C2 appearing and C3 reducing in size after injury. HSC, haematopoietic stem cells, mt, mitochondrial.

### Runx1-positive endocardial cells express smooth muscle genes

Within the individual *runx1-citrine* expressing cell groups, we focussed next on the endocardial/endothelial cells in the wild-type uninjured and wild-type 3 dpci hearts. *citrine/mcherry* double positive endocardial cells showed a highly injury specific upregulation of collagens, for example *collagen 1a1b* (Fig. 5a). Upregulation of collagens in the wound endocardium has been observed before (36, 37), but has not previously been analysed using single cell transcriptomics. Clustering of the *citrine/mcherry* double positive cells showed 2 clusters (C4 and C5) appearing after injury, with upregulation of genes involved in extracellular matrix formation (Fig. 5b-c, Supp Fig. 5a). Interestingly, the endocardial cells of cluster 4 specifically upregulated smooth muscle genes, with high expression of *myh11a, myl6* and *myl9a/b*. *Tagln* (sm22a) was expressed in both clusters 4 and 5, but strongest in cluster 4 (Fig. 5c-d). We were able to confirm this endocardial expression of smooth muscle gene Myh11 on sections (arrowheads Fig. 5e-f, Supp Fig. 5b). These data combined suggests that the *runx1*-positive endocardium is a heterogeneous cell population, with subsets of cells starting to express collagens and/or smooth muscle genes after injury. This combination of high collagen and smooth muscle gene expression is normally a hallmark of myofibroblasts. As the presence of smooth muscle genes in the endocardium has not been reported yet, we looked into this in more detail and expanded our analysis to the *runx1* mutants.

**Fig. 5.**
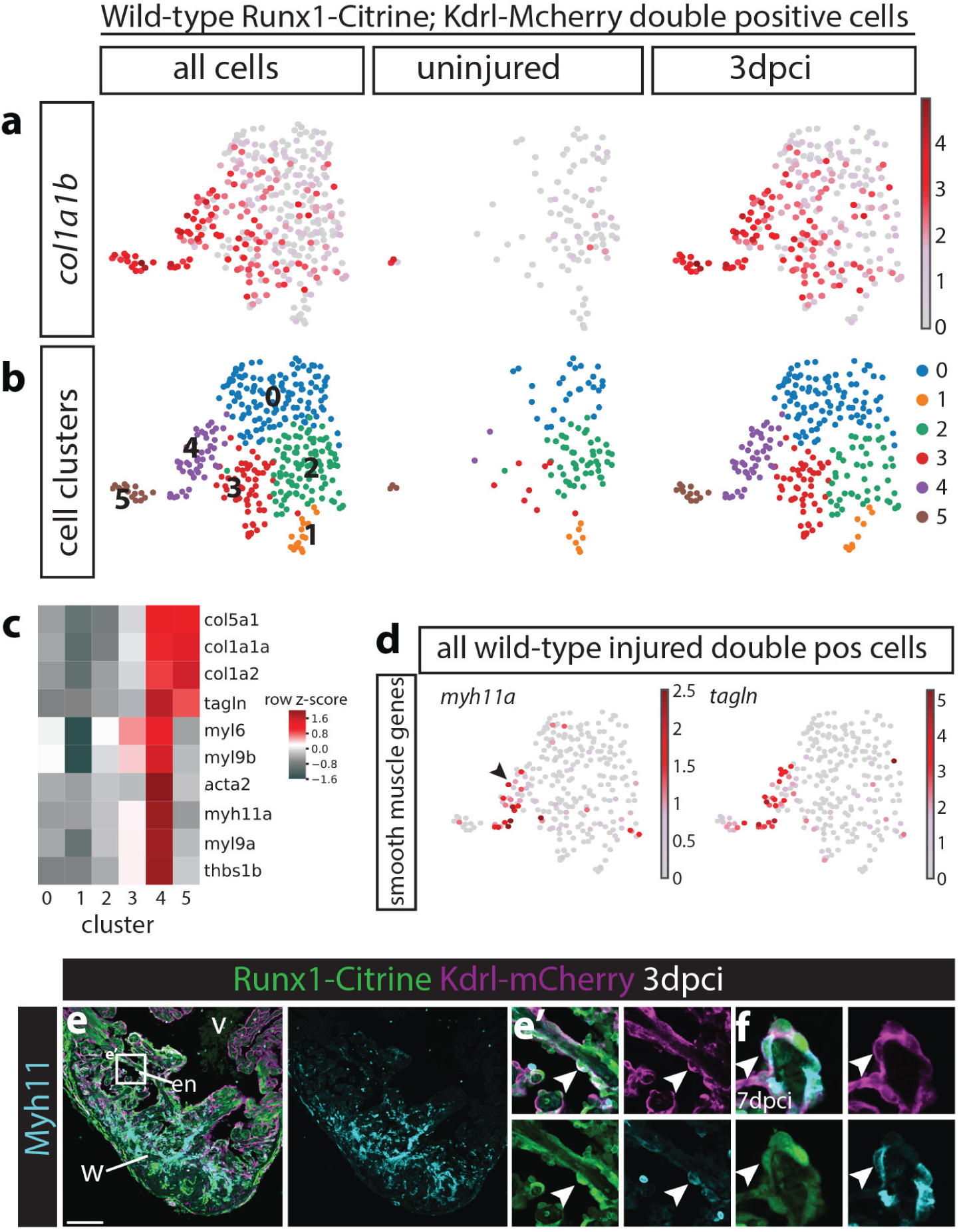
Subset of wild-type double positive cells expresses smooth muscle genes. a-b, all wild-type uninjured and 3dpci *Tg(BAC-Runx1P2:Citrine;kdrl:Hsa*.*HRAS-mCherry)* double positive cells visualised in an UMAP plot. a, expression of *collagen 1a1b* specifically in 3dpci double positive cells. b, cell clustering within the double positive population identifies 6 different cell clusters. c, heatmap showing that both cluster 4 and 5 express high levels of collagens, whereas cluster 4 specifically expresses smooth muscle genes. d, *myh11a* and *tagln* are expressed in cluster 4 after injury. Arrowhead points to *myh11a* expression in cell cluster 4. e-f, immunohistochemistry for Citrine, mCherry and Myh11. Arrowheads point to expression of Myh11 in *Tg(BAC-Runx1P2:Citrine;kdrl:Hsa*.*HRAS-mCherry)* double positive cells at 3 (e-e’) and 7 dpci (f). en, endocardium; v, ventricle; w, wound. Scale bars depict 100 μm.

### Smooth muscle genes are expressed in the endocardium and thrombocytes, but both these cell-types are absent in the *runx1* mutant

We first wanted to see if the lower number of *citrine/mcherry* double positive cells in the mutant obtained a similar smooth muscle profile as the injured wild-type and included the *runx1* mutant to our analysis of double positive cells of the single cell data. While cluster 5 was present, the smooth muscle gene expressing endocardial cluster 4 was completely absent in the *runx1* mutant (arrowheads, Fig. 6a-b) as were the cells most strongly expressing *myh11a* and *tagln* (Fig. 6b). Analysis of Myh11 expression in the wound confirmed that these cells were indeed missing in the mutant, however, the strongly reduced area of cells expressing Myh11 in the mutant (Fig. 6c-d) was too large to be attributed exclusively to endocardial cells. While analysing endocardial Myh11 presence via antibody staining on heart sections, we also noticed Myh11 expression in circulating blood cells in wild-type hearts (arrowheads, Fig. 6e-e’). Myh11 is considered a marker for smooth muscle and myofibroblasts, but in addition to being present in endocardial cells, Myh11 staining also clearly overlapped with strong *itga2b* expressing cells, identifying these cells as thrombocytes (Fig. 6e-e’). The single cell data confirmed that the *itga2b*-positive thrombocyte populations highly express *myh11a* (Fig. 4b-c, Fig. 6f-g) and we confirmed the accumulation of Myh11/*itga2b*-positive thrombocytes in the wound area after injury (arrowheads, Fig. 7a). Also in line with the single cell data, we found that the Myh11-positive thrombocyte population was largely absent in the mutant (Fig. 7a-b’). This means that both smooth muscle gene-expressing cell types, the Myh11-positive endocardium and thrombocytes, are missing in the mutant and raises the question what happens to these cells over time in the wild-type situation and how that relates to the observed differences in fibrin and collagen of the AFOG stained sections (Fig. 2). Surprisingly, at 14dpci, we found that Myh11 is still largely present in the endocardium and thrombocytes in wild-types (arrowheads, Fig. 7c-d’), suggesting that these cells do not lose their cell identity and do not differentiate into pure myofibroblast/smooth muscle cells. This double identity could mean that some of the scar forming cells in fish do not fully differentiate into ‘scar-type’ cells, but remain endocardial or thrombocyte in nature. Interestingly, most collagen (blue) deposition in the wound seen in the AFOG staining (Fig. 3b-c) was observed near the location of the Myh11 expressing endocardial cells and thrombocytes, and to much lesser extend near the epicardium (Fig. 2b-c, 7a). The strong reduction of both the blue AFOG staining and Myh11-positive cells in the mutant suggests, therefore, that most of this collagen is deposited by the Myh11-positive endocardial cells and thrombocytes, indicating that these cell types behave as myofibroblasts in the wild type-setting. Absence of these cells could, therefore, explain the different wound composition as well as the enhanced regeneration in the *runx1* mutant. This is a novel and surprising finding, as the epicardium is considered the main source of myofibroblasts and collagen deposition in the heart (37, 39, 40).

**Fig. 6.**
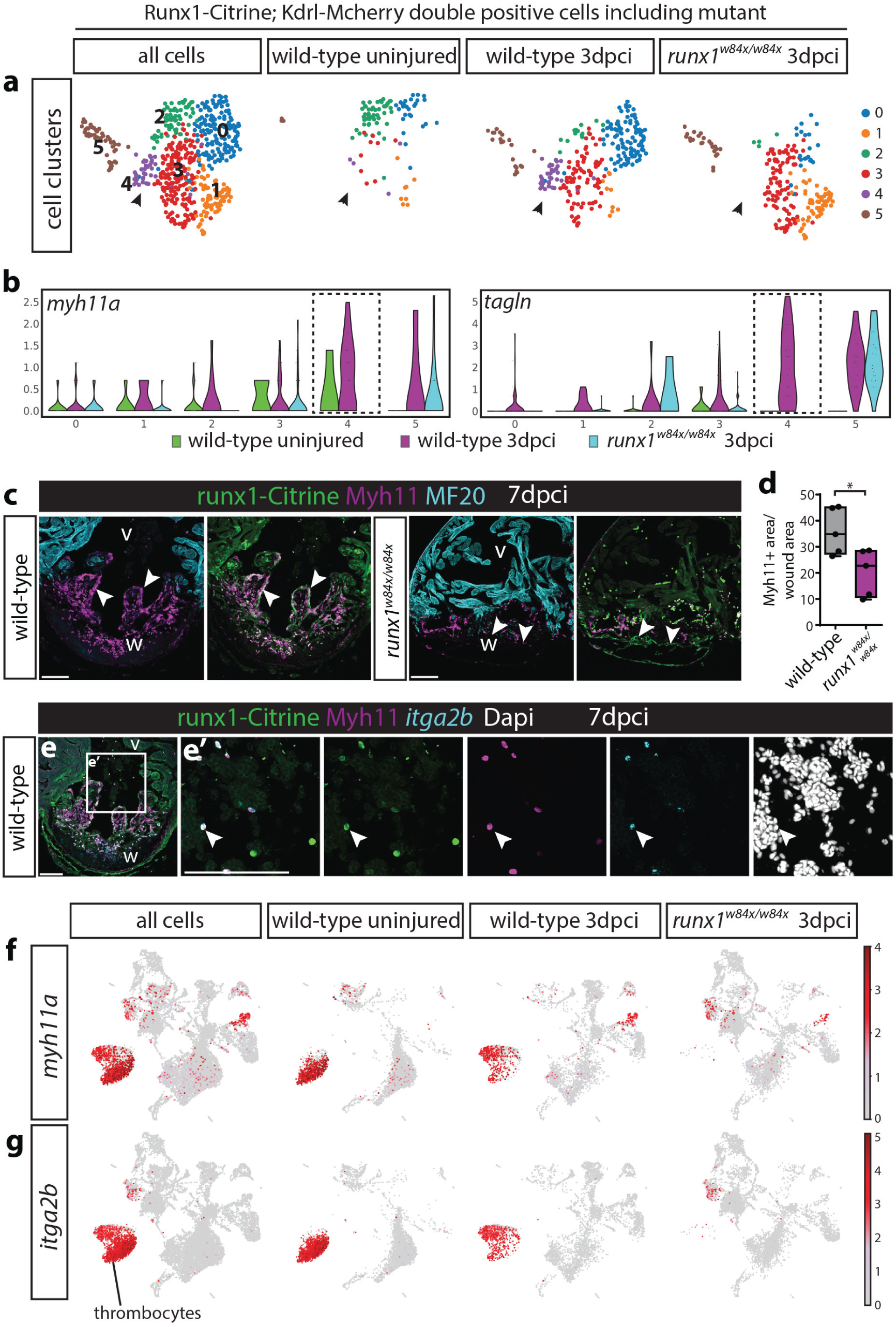
Endocardial and thrombocyte Myh11-positive populations are strongly reduced in the *runx1* mutant. a, UMAP plot of *Tg(BAC-Runx1P2:Citrine;kdrl:Hsa*.*HRAS-mCherry)* double positive cells in the 3dpci *runx1* mutant as well as uninjured and 3dpci wild-type cells. Arrowheads point tot the double positive cluster 4 that is absent in the *runx1* mutant. b, violin plot showing that the absent cluster 4 is the cluster most strongly expressing smooth muscle genes in the wild-type after injury. c, immunohistochemistry for Citrine, Myh11 and MF20 showing reduced staining for Myh11 in the *runx1* mutant wound at 7dcpi compared to the wild-type. Arrowheads point to overlap of Myh11 with Citrine in the endocardium. d, quantification of the area of Myh11 expression in the wound on sections between mutants and wild-types. n=5, unpaired two-tailed t-test. e-e’, *in situ* hybridisation for *itga2b* combined with immunohistochemistry for Citrine and Myh11 with nuclear marker Dapi. Arrowheads point to Myh11 expressing blood cells that express thrombocyte marker *itga2b*. f-g, UMAP plot of all cells confirms expression of *myh11a* in the *itga2b*-positive thrombocyte cluster. v, ventricle; w, wound. Scale bars depict 100 μm. 8 | bioR*χ*iv Koth, Wang and Killen *et al*. | Runx1 acts a negative regulator of heart regeneration

**Fig. 7.**
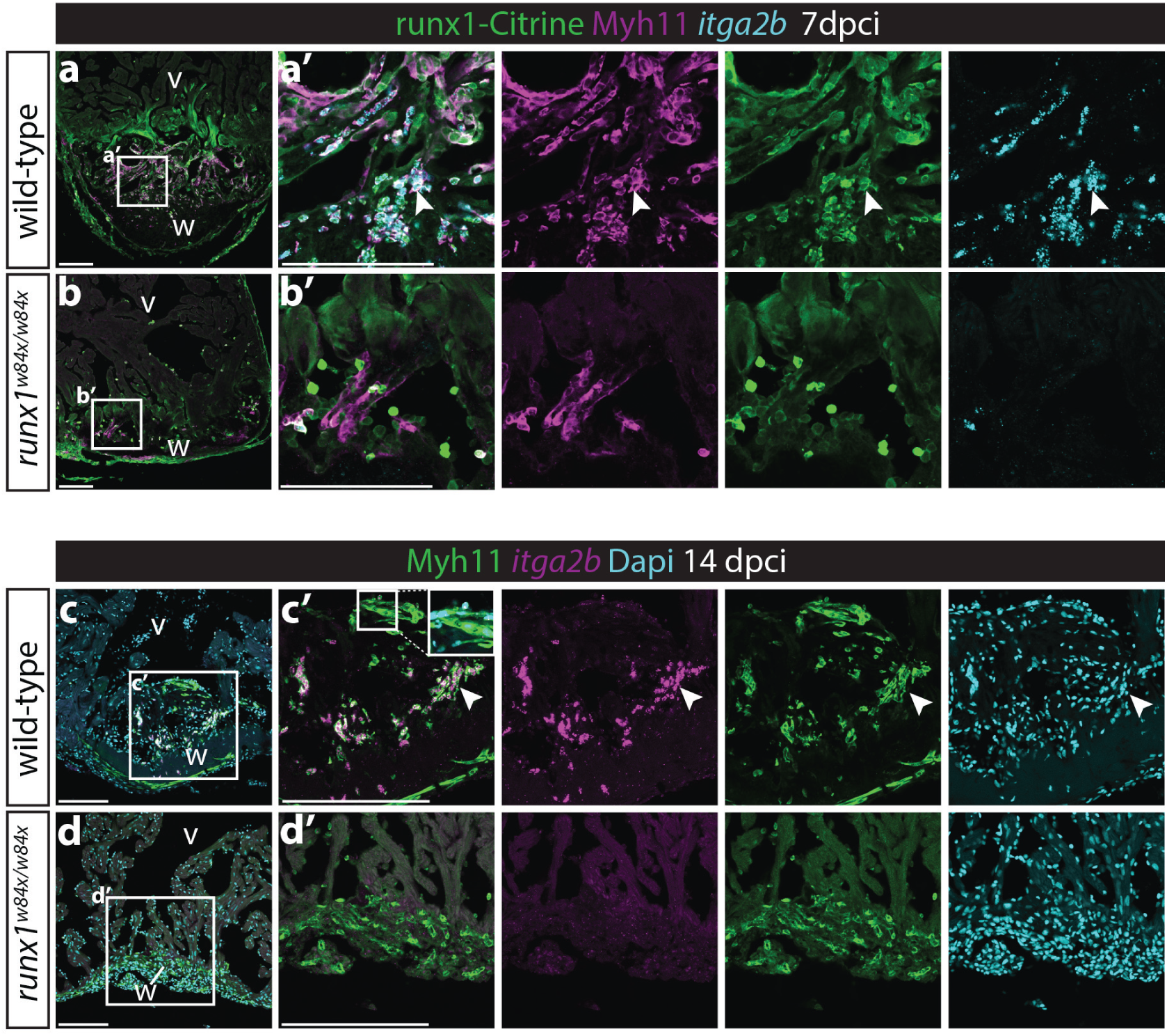
Myh11-positive endocardial cells and thrombocytes retain their double identity at 14dpci. a-b’, *in situ* hybridisation for *itga2b* combined with immunohistochemistry for Citrine and Myh11. Arrowheads point to Myh11-positive *itga2b*-positive thrombocytes present in the wild-type wound, that are largely missing in the *runx1* mutant wound at 7dpci. c-d’, *in situ* hybridisation for *itga2b* combined with immunohistochemistry for Myh11 with nuclear Dapi staining. Both the endocardium (insert) and thrombocytes (arrowheads) still express Myh11 in the wild-type wound at 14dpci, while absent in the mutant wound. v, ventricle; w, wound. Scale bars depict 100 μm.

### Changes in Runx1-Citrine positive epicardial cells after injury

As the epicardium has been shown to contribute myofibroblasts to the wound in zebrafish (37, 40), and we observed activation of Runx1-Citrine in the epicardium, we next compared the epicardial cluster to the myofibroblast/smooth muscle cluster (C8 and C10, Fig. 5b-c, 8a). Cluster 10 is a distinct population of cells specifically appearing after injury that, in addition to the endocardial and thrombocyte populations, strongly expresses both *myh11a* as well as collagens (Fig. 8b). The high levels of expression of smooth muscle genes as well as collagens, suggest that these cells are mainly myofibroblasts, but may also include smooth muscle cells. In the uninjured wild type heart, only a few cells of both populations were positive for Runx1-Citrine (Fig.1a, 8a), but there was a heart-wide activation of Runx1 in the epicardium and myofibroblasts after injury (Fig. 1b-c, 4e, 8a). The epicardial cell cluster specifically expressed a combination of genes known to be epicardial specific, including *tbx18, tcf21* and *wt1a/b*, while *myh11a* expression was specific for the myofibroblast cluster. However, other myofibroblast genes, such as *tagln* and collagens, were present in both the epicardial and myofibroblast clusters (Fig. 8b). Staining on sections confirmed the absence of Myh11, but presence of Tagln in the epicardium (Fig. 8c). The presence of myofibroblast genes in the epicardium might reflect the lineage transition from epicardial cells to myofibroblasts that has been shown before (40). In the *runx1* mutant, similar to the reduction in other *myh11a* expressing cells types, the number of myofibroblasts was much lower. In contrast, the epicardial cell population was slightly larger in the mutant (Fig. 8d) but had reduced levels of both collagen and smooth muscle genes (Fig. 8e). Analysis on sections confirmed that Myh11-positive cells close to the epicardium were less abundant in the mutant compared to the wild-type (arrowheads, Fig. 8f-g). The strong reduction in myofibroblasts in addition to the absence of smooth muscle gene expressing endocardium and thrombocytes, further explains the difference in wound composition between the mutant and wild-type.

**Fig. 8.**
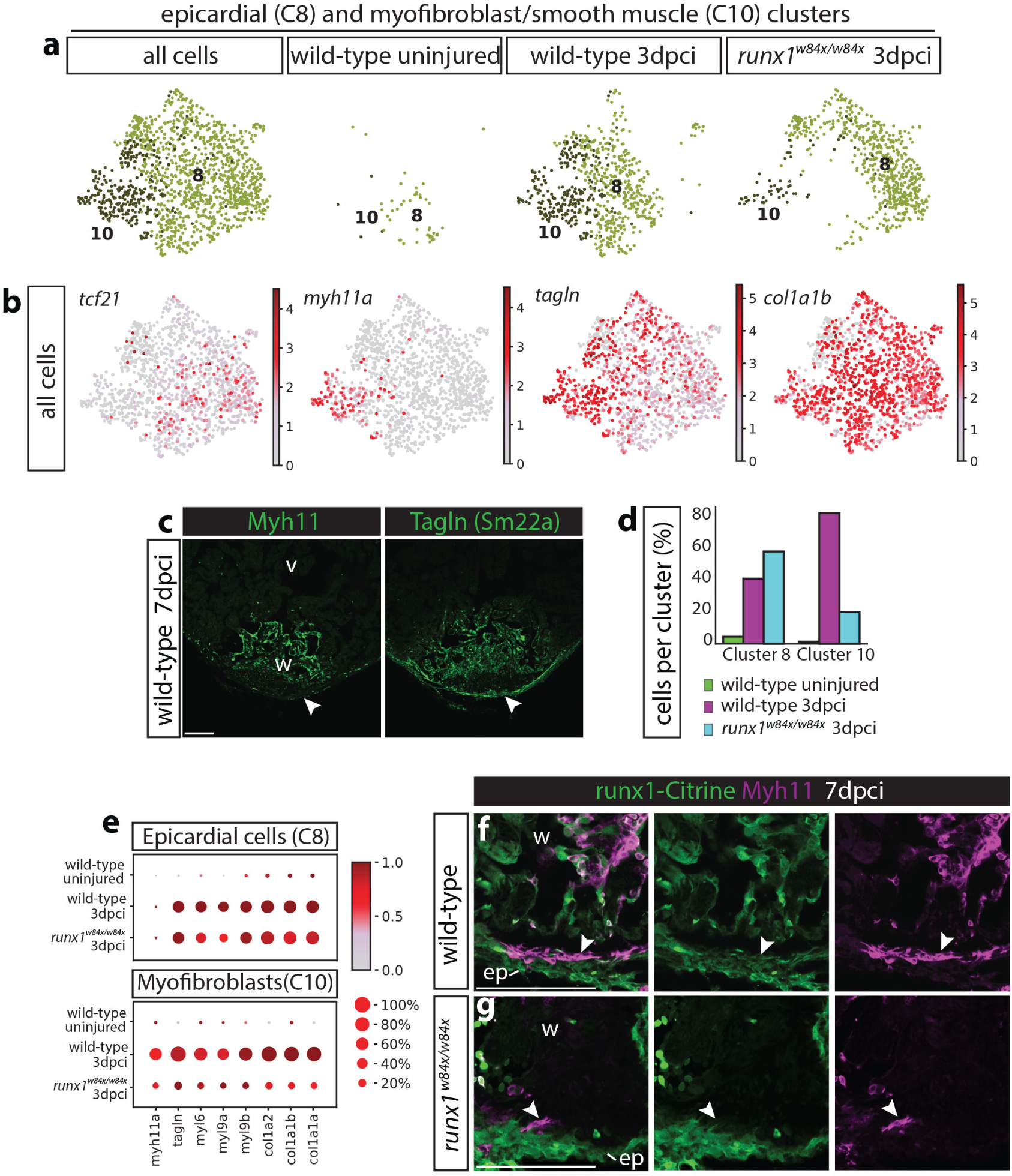
Reduction in myofibroblast numbers in the *runx1* mutant. a, UMAP plot combining cluster 8 and 10 from Figure 4b, showing very few cells in these clusters in the uninjured wild-type, but appearance of both populations after injury in the wild-type and *runx1* mutant. b, UMAP plot from a, indicating expression levels of *tcf21, myh11a, tagln* and *col1a1b*. The epicardial cluster 8 expresses *tcf21*, whereas myofibroblast cluster 10 expresses *myh11a*. Both cluster express *tagln* and *col1a1b*. c, staining of 7dpci sections confirms presence of Tagln and absence of Myh11 in the epicardium. d, single cell data showing numbers of cells per cluster per sample. Increased number of epicardial cells and a reduced number of myofibroblast cells in the runx1 mutant compared the wild-type. e, dotplot showing expression levels of smooth muscle and collagen genes per sample in cluster 8 and 10. Increased cell numbers and expression of myofibroblast genes in both the epicardial and myofibroblast clusters in the injured wild-types compared to the uninjured wild-types. Epicardial myofibroblast gene expression is lower in the runx1 mutant compared to the uninjured wild-type, while the number of myofibroblast cells is strongly reduced in the mutant. f-g, Immunohistochemistry for Citrine and Myh11 on 7dpci sections. Analysis of Myh11 on 7dpci sections confirmed the reduction in myofibroblast cell numbers close to the epicardium (arrowheads). ep, epicardium; ventricle; w, wound. Scale bars depict 100 μm.

### Upregulation of Plasminogen receptor Annexin a2 in the *runx1* mutant myocardium and endocardium

In addition to smooth muscle genes, our single cell sequencing data showed upregulation of a number of genes involved in fibrinolysis in the endocardium after injury, in-cluding *anxa2a* and *s100a10b* (Fig. 9a). Anxa2 is a calcium-dependent phospholipid-binding protein that forms the Annexin A2 hetero-tetramer protein complex together with S100A10 (41) and is an important Plasminogen receptor. Plasminogen is required for dissolving fibrin blood clots and acts as an important protease in tissue remodelling and repair. The Annexin A2 complex was specifically upregulated after injury in the wild-type *runx1-citrine*-positive endocardium, but much stronger in the *runx1-citrine*-positive *runx1* mutant endocardium (Fig. 9a). In contrast, *serpine1* (also called Plasminogen Activator Inhibitor 1), an inhibitor of fibrinolysis, was much stronger upregulated in the injured wild-type endocardium than in the *runx1* mutant (Fig. 9a), suggesting increased fibrinolysis in the *runx1* mutant underlies the reduced amount of fibrin in the wound (Fig. 2). Interestingly, *thbs1b* (Thrombospondin 1b), which is required for thrombocyte and fibrin aggregation, is specifically expressed in the smooth muscle-expressing endothelial population that is missing in the mutant, as a potential link to the absence of thrombocytes and fibrin aggregates in the mutant (arrow-heads, Fig. 9b).

**Fig. 9.**
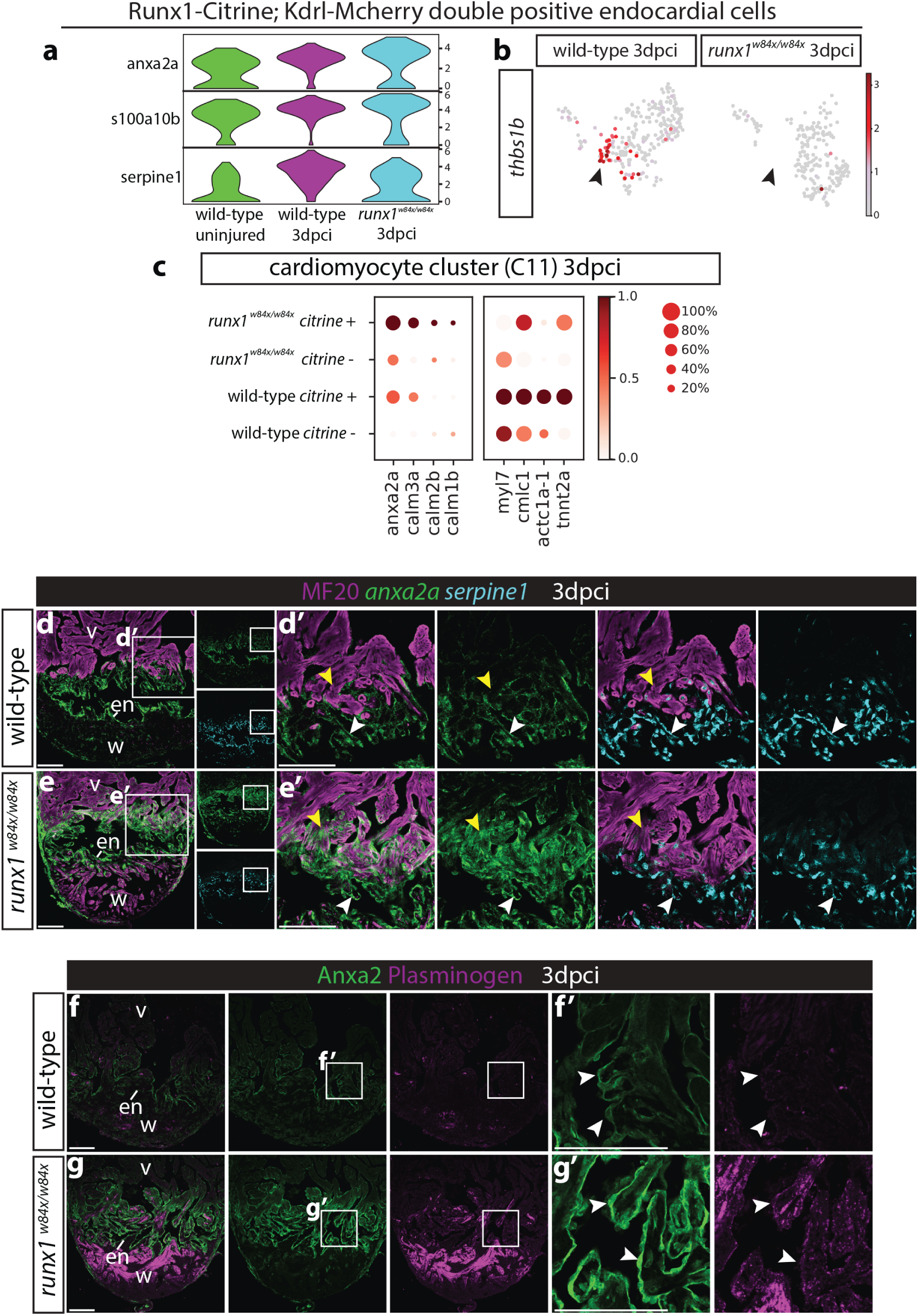
*Runx1* mutant hearts upregulate Anxa2 and Plasminogen. a, violin plots showing upregulation of anx2a and s100a10b in wild-type *runx1/citrine;mcherry/kdrl* double positive cells after injury, with even higher expression in the *runx1* mutant. In contrast, *serpine1* is down regulated in the mutant cells. b, UMAP plot of the *runx1/citrine;mcherry/kdrl* double positive cells showing *thbs1b* expressing cells. Arrowheads point to *thbs1b* expression mainly in cluster 4 from Fig. 6a, which is missing in the *runx1* mutant after injury. c, dotplot showing that *anxa2a, calm1b, calm2b* and *calm3a* are upregulated in the mutant *citrine*-positive myocardium at 3dpci, whereas sarcomere genes are upregulated in the wild-type *citrine*-positive myocardium. d-e’, section *in situ* hybridisation for *anxa2a* and *serpine1* with immunohistochemistry for MF20 shows that *anxa2a* has a similar expression pattern as Runx1-Citrine after injury in the wild-type, but is much higher expressed in the mutant endocardium (white arrowheads) and myocardium (yellow arrowheads). Endocardial *serpine1* expression is lower in the mutant compared to the wild-type (white arrowheads). f-g’, immunohistochemistry for Plasminogen and Anxa2 shows upregulation of Plasminogen in the area where Anxa2 is upregulated in the mutant. en, endocardium; v, ventricle; w, wound. Scale bars depict 100 μm.

In a previous study, cardiomyocyte specific knock-out of *Runx1* in the mouse showed increased sarcoplasmic reticulum calcium content and sarcoplasmic reticulum–mediated calcium release in the myocardium after injury (29). Correspondingly, we found increased expression of calcium-responsive genes in the *citrine*-positive myocardium of our zebrafish *runx1* mutant after injury; including Calmodulins *calm1b, calm2b* and *calm3a* (Fig. 9c, (cluster C11 in Fig. 4b-c). In addition to its expression in the endocardium, we found calcium-dependent *anxa2a* to be the highest upregulated gene in the myocardium in the *runx1* mutant, again suggesting an increased expression of the Plasminogen receptor in the mutant compared to wild-types (Fig. 9c). We confirmed the expression of both *anxa2a* and *serpine1* at 3dpci, which showed that *anxa2a*, as well as Anxa2 protein, has a very similar expression pattern to runx1, however, with much stronger expression in the endocardium (white arrowheads) and myocardium (yellow arrowheads) in the mutant than in the wild-type (Fig. 9d-g’). *Serpine1* expression was reduced in the injured mutant compared with the wild-type, further confirming the single cell data. The increase in *anxa2a* and *s100a10b*, that together convert Plasminogen into Plasmin to increase fibrinolysis, accompanied by the reduction in fibrinolysis inhibitor *serpine1*, all point to substantial differences in Plasmin levels and thus fibrinolysis in the *runx1* mutant. In concordance, Plasminogen was far more abundant in the wound in the mutant (Fig. 9f-g). Therefore, in addition to a lack of fibrin deposition by endocardial cells, thrombocytes and myofibroblasts, *runx1* mutant hearts also show increased fibrinolysis that further explains the observed reduced fibrin presence in the wound and the faster repair in the *runx1* mutant.

### Increased myocardial proliferation and myocardial protection against cryo-injury in the *runx1* mutant

After injury, Runx1-Citrine expression was activated in cardiomyocytes that were in direct contact with the wound (Fig.1). These border-zone cardiomyocytes are known to be highly proliferative and to contribute new cardiomyocytes to the wound (2). We observed increased expression of genes important for sarcomere formation in the *citrine*-positive myocardium in the wild-type hearts, suggesting activation of differentiation of newly generated cardiomyocytes. In the *runx1* mutant, however, this upregulation was reduced (Fig. 9c), suggesting mutant cardiomyocytes were less differentiated, which could result in increased proliferation. To test this, proliferating cells were labelled on sections with Proliferating Cell Nuclear Antigen (PCNA) and myocardial nuclei with Mef2 in addition to the Runx1-Citrine transgene detection. In both wild-type and *runx1* mutant hearts, there was overlapping PCNA and Citrine expression (Fig. 10a-b’). As the down-regulation of sarcomere genes suggested, myocardial proliferation was significantly increased near the wound in mutants compared to the wild-types at all time-points analysed (Fig. 10c). Thus, in addition to its role in the endocardium and thrombocytes, Runx1 expression in the wound border zone cardiomyocytes appears to inhibit myocardial proliferation. Fascinatingly, in the *runx1* mutant wound, we observed a large number of myocardial cells surviving after injury, most notably at 3dpci, which were absent in wild type injured hearts and independent of initial wound size (Fig. 10d-e). These surviving myocardial cells were still present at 7dpci in the mutant, albeit in reduced numbers (Fig. 10d-e), however, they did not express Runx1-Citrine and had lost their normal sarcomere structure (Fig. 10e). This protection of the cardiomyocytes against injury resembles the cardioprotective effect seen in mice, which was found to be link to the improved calcium uptake in the sarcoplasmic reticulum. Taken together, these findings reveal pleiotropic roles for Runx1 in reducing cardiomyocyte proliferation and survival following heart injury.

**Fig. 10.**
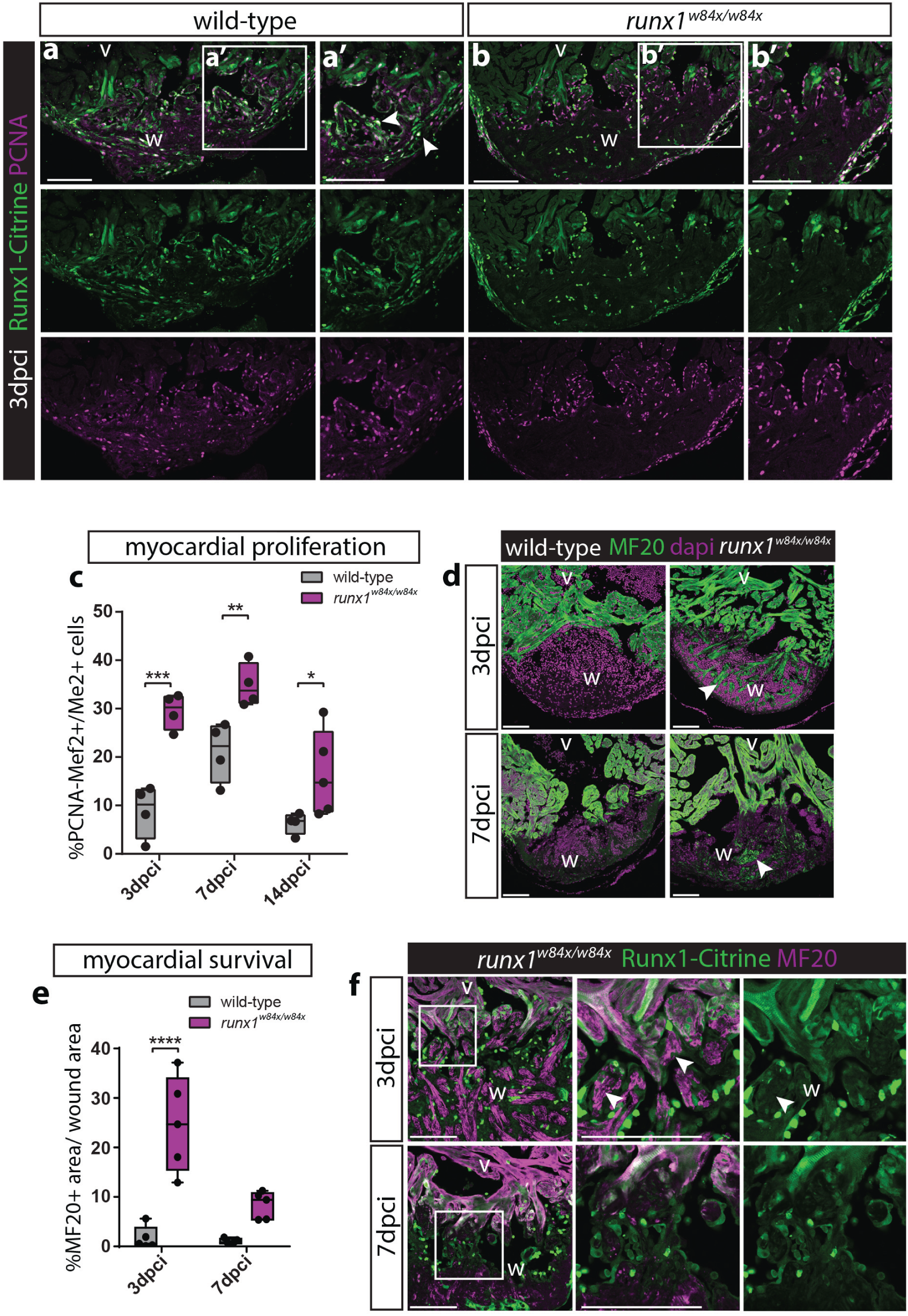
Increased myocardial proliferation and protection in the *runx1* mutant. a-b’, Immunohistochemistry for Citrine and PCNA on 3dpci sections. Runx1-Citrine expression shows large overlap with proliferating cells positive for PCNA in both the wild-type and mutant wound. c, quantification of PCNA-positive proliferating Mef2-positive myocardial cells after injury shows increased myocardial proliferation in the *runx1* mutant at all time-points analysed. n*≥*4, two-way ANOVA with Sidak test. d, Immunohistochemistry for MF20 with nuclear marker Dapi. Arrowheads point to presence of MF20-positive myocardial cells in the wound in the mutant at both 3 and 7 dpci. e, quantification of the MF20-positive area on in the wound on sections between the wild-type and mutant shows increased presence of myocardial cells in the mutant. n=5, two-way ANOVA with Sidak test. f, Immunohistochemistry for Citrine and MF20. Arrowheads point to the surviving MF20-positive cells in the mutant wound that are Runx1-Citrine negative. v, ventricle; w, wound. Scale bars depict 100 μm.

## Discussion

By interrogating Runx1 function on a single cell level in a global *runx1* null mutant, we have exposed *runx1* as an inhibitor of heart repair on many levels. The overarching change we observed in different injury responsive cell-types was a strong reduction of smooth muscle and collagen gene expression in the mutant after injury. The expression of high levels of smooth muscle and collagen genes is a hallmark of myofibroblasts, but we identified two novel cell populations that expressed myofibroblast-like genes after injury: endocardial cells and thrombocytes. The endocardium proximal to the wound has been described before to upregulate collagens (36, 37), but we now show that these cells also upregulate smooth muscle genes such as Myh11 and Tagln (Sm22a). In addition, we found that the Runx1-Citrine positive thrombocytes that make up a large proportion of the wound express Myh11. Surprisingly, we observed most collagen deposition in the wound localised adjacent to the Myh11 expressing endocardial cells and thrombocytes, and to a much lesser extent near the epicardium, despite the fact that the epicardium is considered the main source of myofibroblasts and collagen deposition in the heart (37, 40). This suggests that while these cells retain the characteristics of endocardial cells and thrombocytes, they can function analogously to myofibroblasts. The double identity of these cells may mean that the scar forming cells in fish are more transient and less differentiated compared to fully mature myofibroblasts which in turn may reflect deposition of a less stable, degradable scar compared with the mammalian situation in which myofibroblasts predominate and the scar can persist for many years after MI (42). Runx1 mutants do not have a Myh11 expressing endocardial and thrombocyte population, and have a much smaller number of myofibroblasts. This alone could explain the significant reduction in collagen and fibrin in the wound, however, the *runx1* mutant also shows increased expression of the Plasminogen receptor Annexin A2 as well as Plasminogen itself. Annexin A2 converts Plasminogen into Plasmin, the major fibrinolytic agent that breaks down fibrin in blood clots (41, 43). In addition to a reduction in scar depositing cells, the increased levels of Plasminogen point to faster degradation of deposited scar tissue, allowing improved migration of myocardial cells into the wound to regenerate the heart.

A runx1 enhancer has recently been identified 103kb upstream of runx1 specifically driving expression in the zebrafish wound border myocardium after injury, with the suggestion that this enhancer is a cardiomyocyte regeneration enhancer (CREE) (25). The expectation based on these results was that Runx1 expression in the myocardium near the wound is beneficial for heart regeneration. A positive role for Runx1 was also suggested by its upregulation in hearts treated with Oncostatin M, which has been shown to protect the heart after acute MI (24), as well as in hearts that over-express Erbb2 and show enhanced myocardial proliferation and regeneration (44). High levels of cardiomyocyte Runx1 expression were linked to the reduced differentiation state of cardiomyocytes in these models which facilitates dedifferentiation and increased proliferation leading to improved levels of heart repair. However, this hypothesis was solely based on expression data and not functionally tested. In contrast, our findings demonstrate the opposite in that loss of *runx1* results in enhanced heart regeneration, by increasing myocardial proliferation, increasing myocardial survival, and by altering scar composition as discussed above. This poses the question as to why Runx1 is specifically upregulated during both zebrafish heart regeneration and mammalian heart repair (24, 25, 29, 44) when it seems to function to inhibit key regenerative processes. The answer might lie in the fact that absence of *runx1* causes increased proliferation and upregulation of the Annexin 2a receptor, which is strongly linked to proliferation in cancer (43). Runx1 might function to keep cardiomyocyte proliferation in check and to prevent it from getting out of control during myocardial regeneration. This fits well with the observations that Runx1 acts as a key factor in determining the proliferative and differential state of multiple cell-types (23, 45–48) alongside different functions that correlate with level of Runx1 expression (49, 50), with higher levels shown to result in cell fate transition and differentiation (51).

We observed upregulation of several calcium-dependent genes, including Anxa2, suggesting increased calcium signalling in the *runx1* mutant heart. These data are supported by the results from myocardial specific *Runx1* knock-out mice, which were found to exhibit improved calcium handling in cardiomyocytes, with increased sarcoplasmic reticulum calcium content and sarcoplasmic reticulum–mediated calcium release. The elevated calcium levels protected the cardiomyocytes against adverse remodelling after MI, preserving cardiomyocyte contraction (29). The cardioprotective effect of increased calcium levels can explain the ability of *runx1* mutant cardiomyocytes to survive cryoinjury. Cardiomyocyte-specific absence of *Runx1* in the mouse prevented adverse cardiac remodelling, but did not influence scar size (29). This indicates that absence of *Runx1* function in the myocardium specifically is important for regeneration.

Taken together, our data show that *runx1* functions to regulate scar deposition and degradation, and repress myocardial proliferation and differentiation as well as myocardial survival in the zebrafish heart. The fact that one gene can inhibit multiple aspects of heart regeneration offers the exciting prospect that all these processes can be targeted simultaneously in efforts to achieve human heart repair. Of note, small molecule drugs inhibiting Runx1 have already passed pre-clinical testing in the context of leukaemia treatment (52). Even though the zebrafish is capable of regeneration, the *runx1* mutant shows that this process is not optimal, arising from the need to initiate a fibrotic response for immediate repair and to keep cardiomyocyte proliferation under control during myocardial regeneration. During evolution, adult zebrafish seem to have established a fine balance regulating fibrosis and myocardial proliferation without losing control of cell division; understanding how this balance is maintained may open up novel targets for future therapeutic interventions.

## Methods

### Zebrafish strains and husbandry

All experiments were carried out under appropriate Home Office licenses and in compliance with the revised Animals (Scientific Procedures) Act 1986 in the UK and Directive 2010/63/EU in Europe, and all have been approved by Oxford’s central Committee on Animal Care and Ethical Review (ACER). Adult wild-type (wt) (KCL strain), *Tg(kdrl:Hsa*.*HRAS-mCherry)* (38) and *runx1^W84X^* mutants (30), *TgBAC(runx1P2:Citrine)* (32) were housed in a Techniplast aquarium system [28°C, 14/10 hours light/dark cycle, fed 3x daily with dry food and brine shrimp]. All double transgenic lines on wild-type or mutant background were generated by natural mating.

### Cardiac surgery

All procedures/protocols were carried out in accordance with British Home Office regulations, with respective project licenses held in all contributing labs, approved by Home office inspectors and local representatives. Zebrafish cryoinjury and resection injury of the ventricle were performed as previously reported (53, 54). Briefly, prior to all surgical operations, fish were anaesthetised in MS222 (Sigma). A small incision was made through the thorax and the pericardium using forceps and spring scissors. The abdomen was gently squeezed to expose the ventricle and tissue paper was used to dry the heart. A cryo-probe with a copper filament was cooled in liquid nitrogen and placed on the ventricle surface until thawing was observed. Body wall incisions were not sutured, and after surgery, fish were returned to water and stimulated to breathe by pipetting water over the gills until the fish started swimming again. For sham surgery, the thorax and pericardial sac were opened, but the heart was not injured. All operated fish were kept in individual tanks for the first week after surgery, then fish were combined in larger tanks.

### Tissue Processing

Hearts were extracted and transferred to Ringer solution with heparin sodium salt (50 U/ml) (Ringer composition: 7.2 g NaCl, 0.225g CaCl2.H2O, 0.37g KCl, 0.2175g Na2HPO4.7H2O, 0.02g KH2PO4 at pH 7.4 sterilised by using a 0.22um bottle top filter unit) and rinsed once with Phosphate buffered saline (PBS) or directly isolated in PBS. Hearts were inspected, cleaned and then fixed with 4% Paraformaldehyde (PFA) overnight (O/N) on nutator at room temperature (RT). Samples were rinsed once in PBS, dehydrated into Ethanol (EtOH) at 70%, 80%, 90%, 96% for 2 hours each step and 2×100% for 1 hour each step, followed by a 100% 1-butanol step overnight. The samples were then transferred to paraffin (Paraplast Plus, Sigma-Aldrich P3683) wax at 65°C. Paraffin was refreshed 2x with each step at least 2 hours, prior to mounting in sectioning mould. 10 μm sections were cut using a Leica microtome and section ribbons were stored on black cardboard in shallow stackable plastic trays. Individual sections, evenly distributed throughout the heart e.g. 1 in 6 sections, were selected and mounted on superfrost plus glass slides for histology, histochemistry and RNA labeling (RNAscope).

### Histology

For Acid Fuchsin Orange G-staining (AFOG), dewaxed and water rinsed sections were refixed in Bouin’s solution for 3h at 60°C and then O/N at RT, then washed in ddH2O until sections were white/clear, incubated in aquous 1% phosphomolybdic acid for 5min, rinsed with ddH2O, stained with AFOG solution for 5-10 min, rinsed shortly in ddH2O, dehydrated quickly through 70%, 80%, 90%, 96% and 100% EtOH, quickly cleared in Xylene and mounted in DPX mounting medium under 24 mm x 50 mm cover glasses no 1.5. AFOG solution: Boil 1l ddH2O with 5g of Methyl Blue (Sigma 95290), once cooled - add 10g Orange G (Sigma O7252) and 15g Acid Fuchsin (Sigma F8129) and adjust to pH 1.09 by adding HCL (at 25% concentration).

### Immunohistochemistry

Fluorescent immunohistochemistry was performed as previously described (55). For de-waxing, slides were taken through 2x Xylene 5min, 2x 1min in 100% EtOH 2 min, 1x 96%, 90%, 80% and 70% EtOH, and a final rinse in PBS prior to subsequent staining. De-waxed and rehydrated sections were heated up and then pressure cooked for 4 minutes in antigen unmasking solution (H-3300, Vector Laboratories Inc). Once cooled, sections were placed in PBS before drawing a ring (ImmEdge pen, Vector Laboratories) around the sections. Slides were placed into staining trays providing humidity and blocked using TNB (0.5% TSA blocking reagent, 0.15 M NaCl, 0.1M TRIS-HCL, pH 7.5, NEL702001KT, Perkin Elmer) for 30min at RT. Blocking agent was removed and primary antibody in TNB was added and incubated O/N at RT. Slides were then washed 3x 5 min in PBS before the secondary antibody (Alexa range, Invitrogen) at 1:200 dilution in TNB was added for 2 hours at RT. For some primary antibodies, an additional amplification step was added to enhance the signal using the TSA kit (NEL756001KT, Perkin and Elmer). Instead of an Alexa secondary antibody, a biotinylated secondary antibody was used at 1:200 dilution in TNB for 1/2 hours at RT, followed by 3x 5min washes in PBS prior to 30 min incubation with conjugated Streptavidin-Horse Radish Peroxidase (Vector Laboratories, SA-5004) and subsequent 3x 5 min washes in PBS. Then either Fluorescein or Tetramethylrhodamine (in DMSO) diluted at 1:100 in amplification buffer was added to the sections for 3 minutes, followed by 3x 5 min washes with PBS and staining with DAPI (2.5μg/ml, Sigma). Slides were mounted in Mowiol 4-88 (Applichem) and slides incubated at 37° C O/N in the dark. The following primary antibodies were used: chicken polyclonal against Green Fluorescent Protein (GFP, 1:200, Aves Lab, GFP-1020), mouse monoclonal against mCherry (clone 1C51, 1:200, Abcam, ab125096), Proliferating Cell Nuclear Antigen (PCNA, clone PC10, 1:200, Dako Cytomation, M0879), Myosin Heavy Chain (MF20, 1:50, HSHB AB-2147781) and Plasminogen (Plg, 1:200, RD systems, MAB1939). Rabbit polyclonal against Lysozyme (LyC, 1:200, Anaspec, AS-55633), ETS Transcription Factor ERG (ERG, 1:200, Abcam, ab110639), Myocyte Enhancer Factor 2 (Mef2 C-21, 1:200, Santa Cruz, sc-313), smooth muscle Myosin heavy chain 11 (Myh11, 1:200, Abcam, ab125884) and Annexin A2 (Anxa2, 1:200, Invitrogen, PA5-14317). For double labelling with RNAscope probes, RNAscope was performed first and then processed for immunohistochemistry as described above, starting from the blocking step. Images were processed in ImageJ to generate magenta and green color combinations.

### RNAscope In Situ Hybridisation

RNAscope® (Advanced Cell Diagnostics, Hayward, CA) (34) was performed on 10μm thick paraffin sections, processed as described above. Sections were baked at 60° C for one hour before deparaffinisation using 2x 5 min Xylene steps followed by 2x 2 minutes 10% EtOH. The slides were air-dried followed by incubation in RNAscope Hydrogen Peroxide (H202) for 15 minutes before washing in MilliQ. The slides were then boiled at 98-102° C for 15 minutes in 1x RNAscope® Target Retrieval solution, placed in 100% EtOH for 3 minutes and air-dried. The sections were the incubated with RNAScope Protease III in a Hybez oven at 40°C for 15 min, washed in MilliQ 2x 2 minutes, followed by incubation with the different RNAScope probes for 2 hours at 40°C. The RNAscope Multiplex Fluorescent Detection Reagents v2 and the TSA Plus Cyanine 3 and 5 fluorophore (Perkin Elmer, NEL744001KT) were applied according to the manufactures instructions. The slides were further processed for immunohistochemistry or mounted in Mowiol 4-88. Advanced Cell Diagnostics designed the probes. Probes used were Dr-tcf21-C2 (485341-C2), Dr-itga2b-C2 (555601-C2), Dr-runx1 (433351), Dr-myb-C3 (558291-C3), Dr-gata2b-C2 (551191-C2), Dr-anxa2a (587021) and Dr-serpine1 (551171-C3).

### Image acquisition and data analysis

Images were acquired using either a Zeiss LSM880 or Olympus FV3000 confocal. Images were processed in FIJI/ ImageJ to generate a magenta/green/cyan/grey color scheme. For all quantifications on sections, individual sections were mounted, evenly distributed throughout the heart e.g. 1 in 6 sections, to reduce the number of hearts needed and guarantee even coverage of the entire heart. Using Fiji/ image J, myocardial regeneration was then quantified by measuring the perimeter of the ventricle of each heart section of AFOG-stained hearts and the length of open compact myocardium. The biggest open myocardium length and ventricle perimeter measurement from each fish was then taken and the open myocardium length was divided by the ventricle perimeter and multiplied by 100 to give the percentage of the myocardium that was still open. Scar area and ventricle area were also measured for each section and again the biggest measurement of each for each fish was used to calculate the size of the scar.

This was done by dividing the scar area by the ventricle size, then multiplying by 100 to give a percentage. This was performed for 5 of each fish type and time point. For analysis of the colour of the wound area on the sections stained with AFOG, we split the colour photo of the wound area up into a red, green and blue channel. The images were then thresholded using the same settings for all hearts for the red and blue channel. The orange area was determined by subtracting the red and blue areas from the total area. Myocardial proliferation was assessed by using Mef2, a nuclear myocardial marker, and proliferating cell nuclear antigen (PCNA), a nuclear marker of proliferation, on antibody stained sections. The border zone region was established as the cardiomyocytes closest to the wound in the healthy myocardial tissue. The number of Mef2+ nuclei was counted and the number of PCNA Mef2 double+ nuclei counted and their percentage calculated for at least 3 sections per fish, for four fish per condition.

### Heart processing for FACS

Freshly isolated hearts were placed in chilled Hanks balanced buffered saline (HBBS), atria and bulbus arteriosus were removed and the ventricle was cut into several pieces using fine forceps and ophthalmic scissors (FST, 15009-08). The following digestion procedure was adapted from (56): Pieces were transferred to a 2 ml tube, rinsed with HBBS, 1 ml (up to 10 hearts) of digestion mix (0.13 U/ml Liberase DH (Roche) and 1% sheep serum in HBBS) was added and incubated at 32° C and 80-100rpm rotation/agitation. Every 10-15 min supernatant was collected and placed on ice and new digestion mix was added and pieces and solution where gently pipetted 5-10 times to aid break up. Once all tissue was resolved (1h) all collected suspensions were spun at 300g for 10min, supernatant was removed and replaced with 1% fetal bovine serum in HBBS, suspensions were then combined and placed on ice. Prior to FACS (MoFlo Asterios, Beckman and Coulter) cells were stained with Dapi to gate for dead cells.

### Single cell sequencing

Cells from 20 hearts per sample from isolated uninjured WT and 3 dpi injured WT and *runx1^W84X/W84X^* ventricles were FACS sorted and populations of single and double positive cells were isolated separately. The single and double positive cells were then mixed, so that the samples were 1/3 Citrine positive, 1/3 double positive and 1/3 mCherry positive. Cells were washed in PBS with 0.04% BSA and re-suspended before loading 12,000-12,500 cells onto each channel of the Chromium 10x Genomics platform to capture single cells in droplets. Library generation for 10x Genomics v2 chemistry was performed following the Chromium Single Cell 3 Reagents Kits User Guide: CG00052. Quantification of cDNA was performed using Qubit dsDNA HS Assay Kit (Life Technologies Q32851) and high-sensitivity DNA tape-station (Agilent. 5067-5584). Quantification of library construction was performed using Qubit dsDNA HS Assay Kit (Life Technologies Q32851) and high-sensitivity DNA tape-station (Agilent. 5067-5584). Libraries were sequenced on Illumina HiSeq4000 platform to achieve a minimum of 20,000 reads per cell.

### Data analysis

#### Alignment

The count function in 10x Genomics Cellranger software (v2.1.1) was used for sample demultiplexing, barcodes processing and gene counting with – chemistry=threeprime. (https://support.10xgenomics.com/single-cell-gene-expression/software/pipelines/latest/installation). The Danio_rerio.GRCz11 (release 94) version of Zebrafish genome and gene annotation files used for alignment were downloaded from the Ensembl database. (ensembl.org) We added the sequences for the mCherry-plasmid and Citrine-plasmid. The mCherry-plasmid had three sequence contents: mCherry, the mCherry-plasmid-backbone and mCherry-polyA. The Citrine-plasmid had seven sequence contents: Citrine-3x-HA-tag, Citrine-BirA, Citrine-Tav-2a, Citrine, Citrine-polyA, Citrine-Frt1 and the remaining plasmid backbone sequences (Citrine-Remaining). The new reference with the additional sequences was built using the mkgtf function in Cellranger. In order to eliminate potential reads that were aligned to both plasmids, only uniquely mappable reads were considered for STAR alignments with an additional parameter –’outFilterMultimapNmax’, ‘1’, added in the reference.py file.

#### Quality control

Quality control was performed from the raw counts with all barcodes. Firstly, cells with less than 100 genes expressed were removed. 4720, 4754 and 6268 cells passed for the wild-type uninjured, wild-type 3dpci and *runx1^W84X/W84X^* 3dpci samples respectively. Secondly, doublets were filtered using the scrublet (57) package in Python. Cells with doublet score larger than 0.3, 0.27 and 0.38 for the wild-type uninjured, wild-type 3dpci and textitrunx1^W84X/W84X^ 3dpci samples were removed respectively. This excluded 55 wild-type uninjured cells, 58 wild-type 3dpci cells and 214 *runx1^W84X/W84X^*3dpci cells. After quality control 4665 wild-type uninjured cells, 4696 wild-type 3dpci cells and 6054 *runx1^W84X/W84X^* 3dpci cells were retained. Non-expressed genes were removed and cells were normalized to 10,000 for each cell and log-transformed.

#### Defining cell types based on marker genes

mCherry positive (mChr+) cells were defined as a union of cells that have at least 1 unique molecular identifier (UMI) count in either mCherry, mCherry-plasmid-backbone or mCherry-polyA. Citrine positive (Cit+) cells were defined as a union of cells that have at least 1 UMI in either citrine, citrine-polyA or citrine-remaining. If the cell had at least 1 UMI for kdrl/runx1 gene, then the cell was labelled as kdrl positive (kdrl+) or runx1 positive (runx1+). kdrl+mChr+ cells were defined as the cells that were either kdrl+ or mChr+. Runx1+cit+ cells were defined as the cells that are either runx1+ or cit+. Finally, the double positive (double+) cells were defined as cells that are both kdrl+mChr+ and runx1+cit+. The number of cells for each cell type is summarised in the table below:

For extracting subsets of cells, including cardiomyocytes (255 cells), double+ cells (561 cells) and double+ without *runx1^W84X/W84X^* cells (361 cells), the same quality control pipeline was applied.

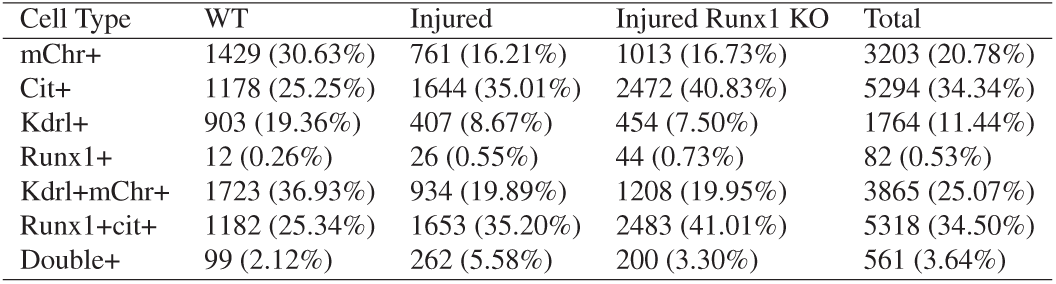

#### Selection of highly variable genes

Highly variable genes (HVGs) were selected following the Seurat (58) method with parameters: min_mean=0.0125, max_mean=4 and min_disp=0.5. 4663, 2341, 3484, 3377, 3854 HVGs were selected for all cells, CMs, double+ cells, double+ no Injured *runx1^W84X/W84X^* cells and epicardial/smooth muscle cells. Cells were then log-transformed. The effects of total number of counts and the percentage mitochondrial genes were regressed out and each gene was scaled so that it was zero-centred.

#### Visualization and Clustering

For UMAP visualisation (Uniform Manifold Approximation and Projection), firstly, a k=10 nearest neighbour graph was calculated on the first 50 principle components of the PCA based on HVGs using the neighbours function in Scanpy (59). Then the UMAP was calculated based on this k-nearest-neighbour graph using the UMAP function in Scanpy. The sub cell populations were determined by Louvain clustering with resolution 1, 0.5, 0.6, 0.7, 0.3 for all cells, cardiomyocytes, double+ cells, double+ without *runx1^W84X/W84X^* cells and epicardial/smooth muscle cells respectively. In total, 26 clusters were defined in all cells, 3 clusters for CMs, 5 clusters for both double+ cells and double+ without *runx1^W84X/W84X^* and 4 clusters for epicardial/smooth muscle cells.

#### Differential expression and Gene Ontology Annotation

Differential expression (DE) analysis was performed using rank_genes_groups in Scanpy with the ‘t-test_overestim_var’ method that overestimates variance of each group. The p values were corrected by the ‘benjamini-hochberg’ (BH) method to account for the multiple comparisons problem. Gene ontology (GO) information was downloaded from the ZFIN database. (https://zfin.org/downloads) Only GO terms with more than 5 genes and less than 500 genes were considered. The enriched GO terms were calculated by a hypergeometric test on the top 50 genes using phyper function in R. Then the p values were then corrected by the ‘benjaminihochberg’ (BH) method using p.adjust function in R.

#### Data plotting

The violin and heatmap plots were made using seaborn and matplotlib modules in Python, and the dotplots using the dot-plot function in Scanpy. The dotplots of the epicardial and the myofibroblast clusters (Fig.8e.) were coloured by the scaled mean expression value by dividing its maximum. The size of dots indicates the number of cells expressing the selected genes for each group. This number was scaled by dividing its maximum. The dotplot of the CM cluster (Fig.9c.) was coloured by the scaled mean expression value for each group, calculated by substracting the minimum and dividing each by its maximum. The dot size was represented as the fraction of cells expressing the selected genes.

### Statistical analysis

The number of samples (n) used in each experiment is shown in the legends and recorded in detail below. Appropriate sample sizes were computed when the study was being designed and no data was excluded. ANOVA tests were applied when normality and equal variance tests were passed. Measurements and counts were performed blinded. Results are expressed as mean ± SEM. (* for P < 0.05, * * for P< 0.01, * * * for P< 0.001 and * * * * for P<0.0001). Statistical analysis was performed in GraphPad Prism 6 for Windows, GraphPad Software, La Jolla California USA, www.graphpad.com. Fig. 2f: Two-way ANOVA with Sidak test, all time points n=5 wild-types, n=5 runx1 mutants. Comparing wild-type versus mutant per time point. 3dpci p=0.1951, 7dpci p=0.9486, 14dpci p=0.0034, 30dpci p>0.9999, 70dpci p=0.9973. Fig. 2g: Two-way ANOVA with Sidak test, all time points n=5 wild-types, n=5 runx1 mutants. Comparing wild-type versus mutant per time point. 3dpci p=0.0222, 7dpci p=0.7108, 14dpci p=0.1220, 30dpci p>0.3548, 70dpci p=0.7223. Fig. 2i-k: Two-way ANOVA with Sidak test, n=5 wild-types, n=5 runx1 mutants. Orange p=0.0001, red p=0.0220, blue p=0.1312. Fig. 3e: Two-way ANOVA with Sidak test. Uninjured, sham, 1dpci and 14dpi n=4. 3 and 7dpci n=5. Comparing time points within wound or ventricle. All ventricle comparisons p>0.9999. Wound: uninjured vs sham p=0.9954, uninjured vs 1dpci p=0.0025, uninjured vs 3dpci p=0.0005, uninjured vs 7dpci p=0.0970, uninjured vs 14dpci p>0.9999, sham vs 1dpci p=0.0501, sham vs 3dpci p=0.0144, sham vs 7dpci p=0.7469, sham vs 14dpci p>0.9999, 1dpci vs 3dpci p>0.9999, 1dpci vs 7dpci p=0.8847, 1dpci vs 14dpci p=0.0118, 3dpci vs 7dpci p=0.6164, 3dpci vs 14dpci 0.0029, 7dpci vs 14dpci p=0.3351. Fig. 6d: Unpaired, two-tailed, equal variance t-test. n=5 wild-types, n=5 runx1 mutants, p=0,0239. Fig. 10c: Two-way ANOVA with Sidak test, all time points n=5 wild-types, n=5 runx1 mutants. 3dpci p=0.0002, 7dcpi p=0.0095, 14dpci p=0.0452. Fig. 10e: Two-way ANOVA with Sidak test, all time points n=5 wild-types, n=5 runx1 mutants. 3dpci p<0.0001, 7dpci p=0.0775.

## Acknowledgements

We thank the Oxford Genomics Centre at the Wellcome Centre for Human Genetics (funded by Wellcome Trust grant reference 203141/Z/16/Z) for the generation and initial processing of the sequencing data. Paul Liu for providing the runx1W84X line, Didier Stainier for the kdrl-mCherry line, Nadia Mercader and Caroline Pellet-Many for training in the cardiac injury models, Michal Maj and Line Eriksen from the Flow Cytometry facility at the Dunn School of Pathology for help in FACS sorting. Ricardo Henriques for the BioRxiv template. Oxford’s BMS teams for zebrafish husbandry support. This work was supported by a BHF non-clinical PhD fellowship FS/14/73/31107 (A.K and M.T.M.M.), BHF project grants PG/15/111/31939 (M.T.M.M.) and (PG/14/39/30865) (J.K. and R.K.P), by the European Research Council (ERC) under the European Union’s Horizon 2020 research and innovation program (grant agreement n° 715895, CAVEHEART, ERC-2016-STG, M.T.M.M.), by a BHF chair award (CH/11/1/28798) and program grant (RG/08/003/25264) (P.R.R) and a MRC-MHU grant (4050189188) (R.K.P.). This work was also supported by the BHF Centre of Regenerative Medicine (RM/13/3/30159) and the BHF Centre of Research Excellence Oxford (RE/13/1/30181). XW and BG were supported by Wellcome, the MRC, CRUK, NIH-NIDDK and core funding from the Wellcome and MRC to the Cambridge Stem Cell Institute. The imaging facilities used for this study are supported by the MRC via the WIMM Strategic Alliance (G0902418), the MHU (MC_UU_12009), the HIU (MC_UU_12010), the Wolfson Foundation (18272) and the Wellcome Trust (Micron 107457/Z/15Z) grants.

---

**Fig. S1.**
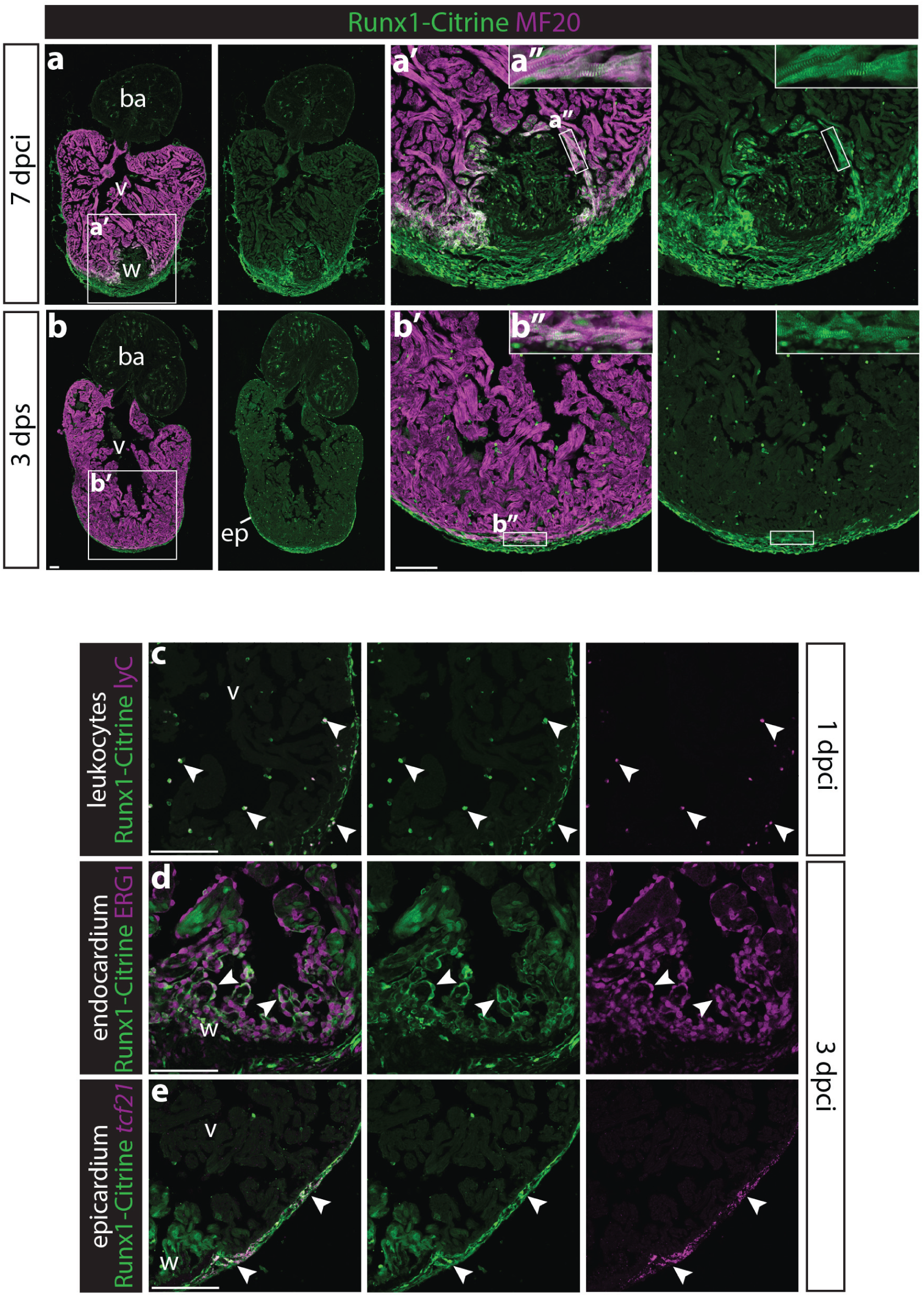
Runx1-Citrine becomes strongly expressed in the heart after cryo-injury. a-b”, immunohistochemistry for Runx1-Citrine (GFP antibody) and myocardial marker MF20 at 7dpci as well as in the sham heart. a-a’, at 7 dpci, the epicardium, endocardium and other wound cells were positive for Citrine. Also the myocardium in the border zone next to the wound was highly Citrine-positive (a”). b-b”, touching the heart with the probe without freezing cells and isolating the heart 3 days later (days post sham, dps) also initiates a response, with Citrine expression in the epicardium and myocardium. c-e, immunohistochemistry for Citrine, LyC, ERG1 and *in situ* hybridisation for *tcf21*. Arrowheads point to overlap of Runx1-Citrine with leukocyte marker lyC at 1dpci (c), and with endocardial marker ERG1 (d) and epicardial marker *tcf21* at 3dpci (e). a, atrium; ba, bulbus arteriosus; ep, epicardium; v, ventricle; w, wound. Scale bars depict 100 μm.

**Fig. S2.**
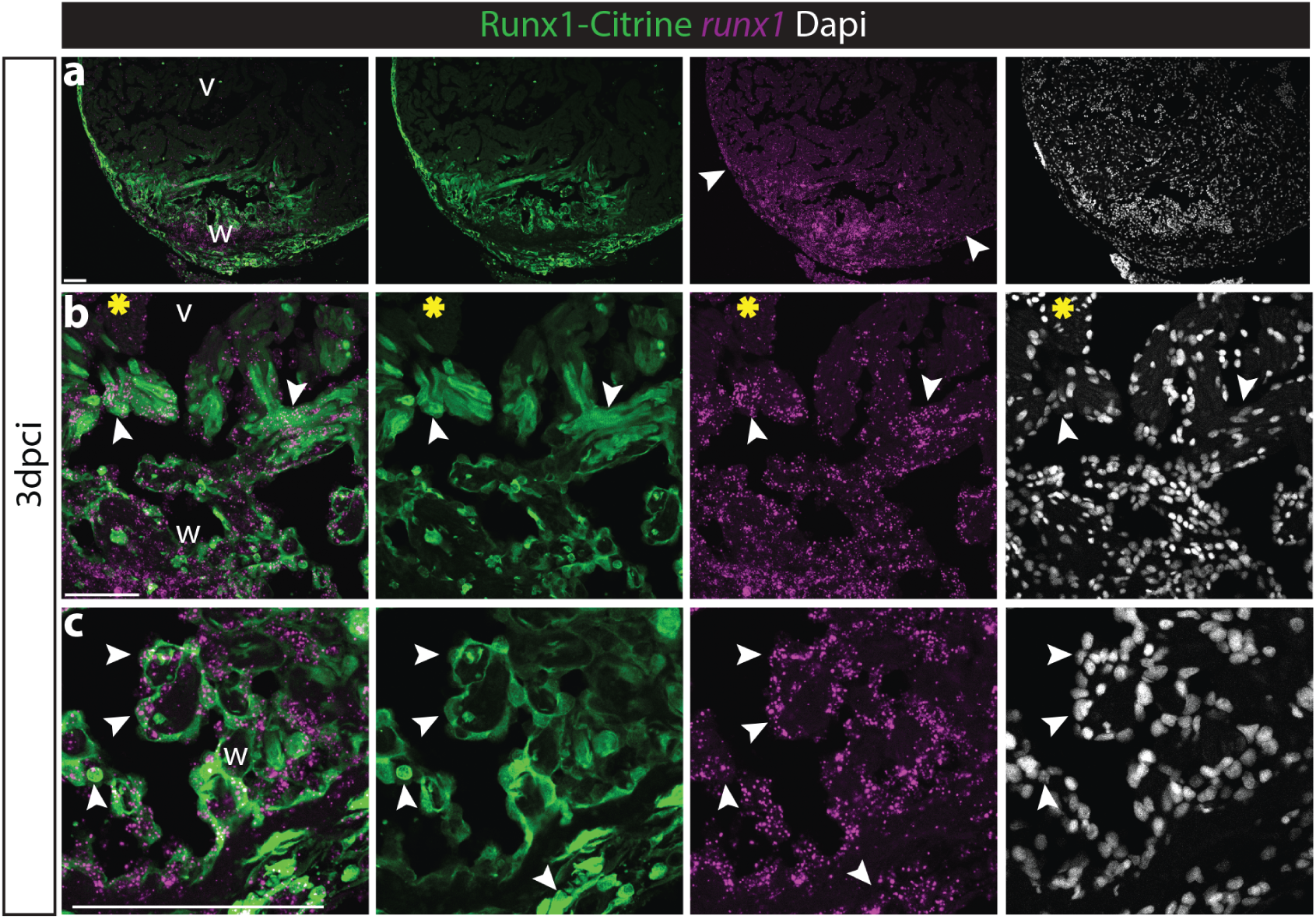
Runx1-Citrine expression recapitulates endogenous *runx1* expression. a-c, immunohistochemistry for Citrine combined with *in situ* hybridisation for *runx1* showing overlap of Runx1-Citrine with runx1 mRNA in wild-type hearts. a, arrowheads point to double positive cells in the epicardium. b, arrowheads point to double positive cells in the myocardium, where the yellow asterisks point to double negative myocardium. c, arrowheads point to double positive cells in the endocardium. v, ventricle; w, wound. Scale bars depict 100 μm.

**Fig. S3.**
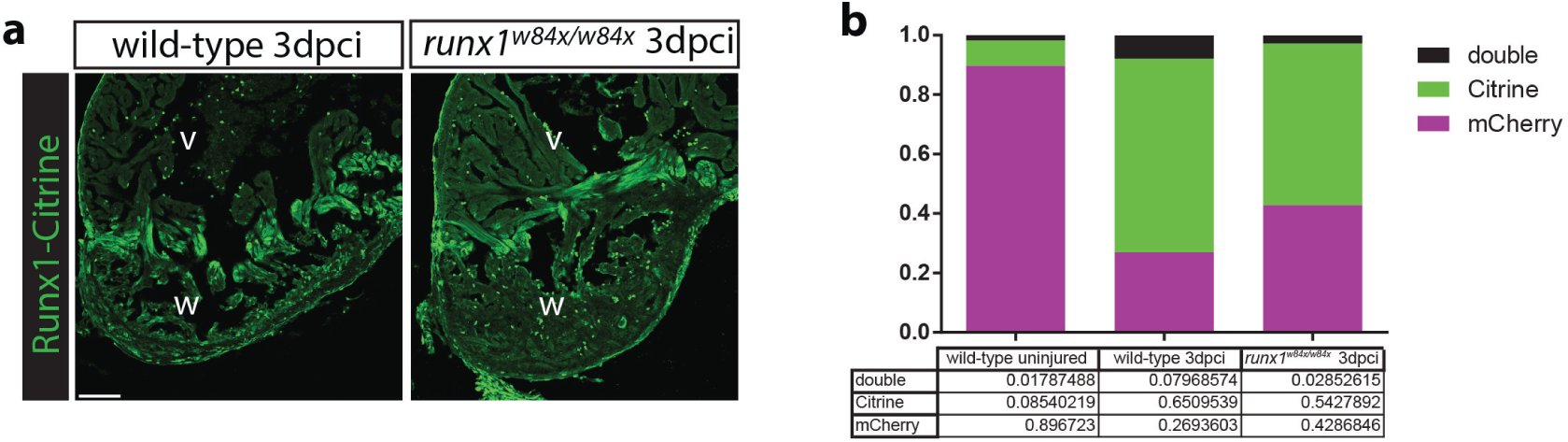
Reduced number of Citrine-positive endocardial cells in the *runx1* mutant. a, immunohistochemistry for Citrine at 3dpci. Similar expression of the Runx1-Citrine protein between wild-type and *runx1* mutant hearts after injury. b, FACS sorting for Runx1-Citrine and kdrl-mCherry shows an increase in double positive cells after injury in the wild-type compared to the uninjured wild-type hearts, whereas there is a reduction in the number of double positive cells in the injured *runx1* mutant compared to the injured wild-types. v, ventricle; w, wound. Scale bars depict 100 μm.

**Fig. S4.**
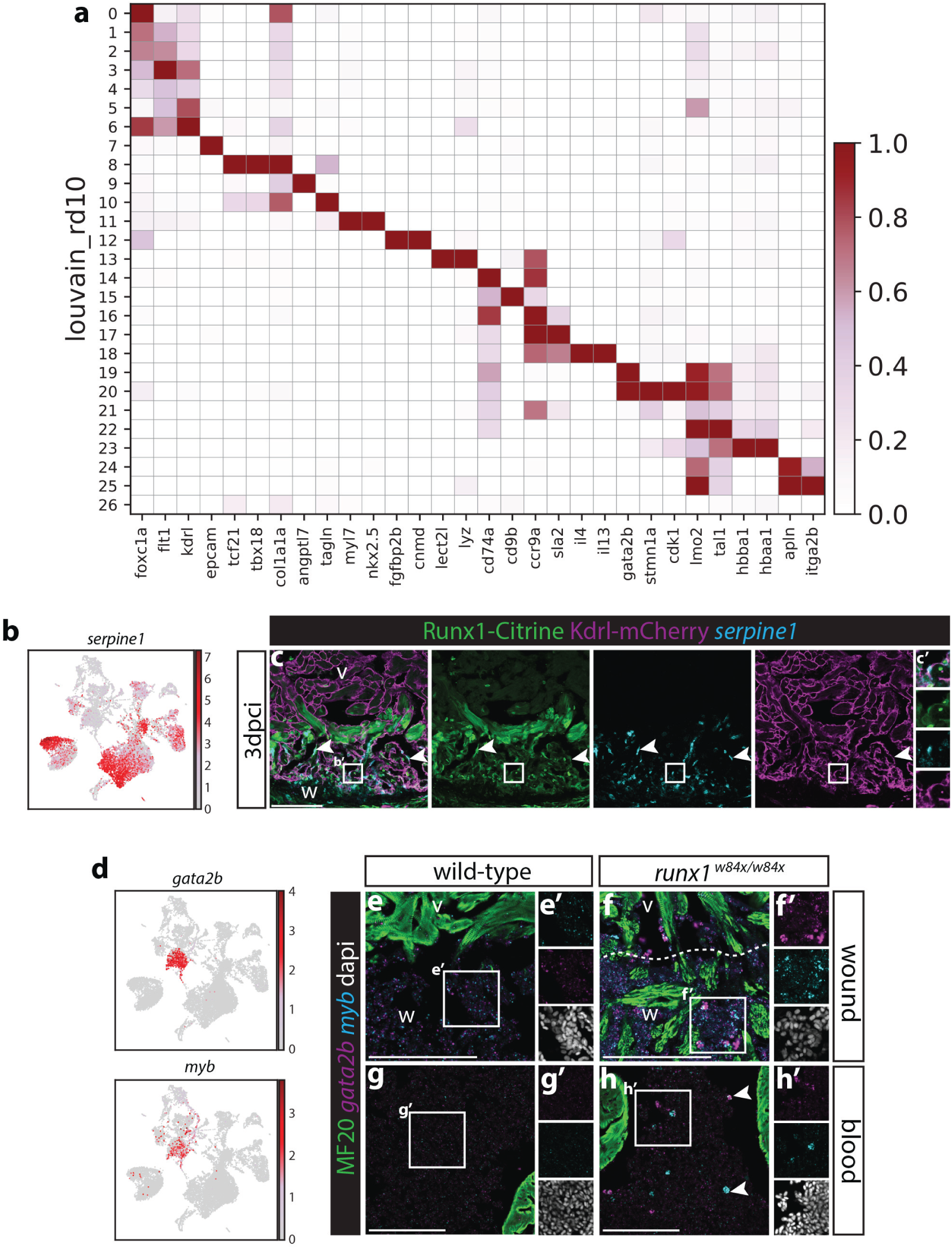
Serpine1 expression overlaps with Runx1-Citrine after injury. a, heatmap showing example genes used to determine the identity of the different cell clusters. b, UMAP plot of all cells showing expression of *serpine1*. c-c’, immunohistochemistry for Citrine and mCherry combined with *in situ* hybridisation for *serpine1*. Analysis of *serpine1* on 3dpci sections shows a largely overlapping expression pattern to Runx1-Citrine in the endocardium (arrowheads and insert c’). d, UMAP plot of all cells show that runx1 mutant specific blood cell populations have high levels of expression of *gata2b* and *myb*. e-h’, immunohistochemistry for MF20 combined with in situ hybridisation for *gata2b* and *myb*. Inserts and arrowheads point to *runx1* mutant specific *gata2b* and *myb* expression in blood cells, both in the wound (e-f’) and blood (g-h’). v, ventricle; w, wound. Scale bars depict 100 μm.

**Fig. S5.**
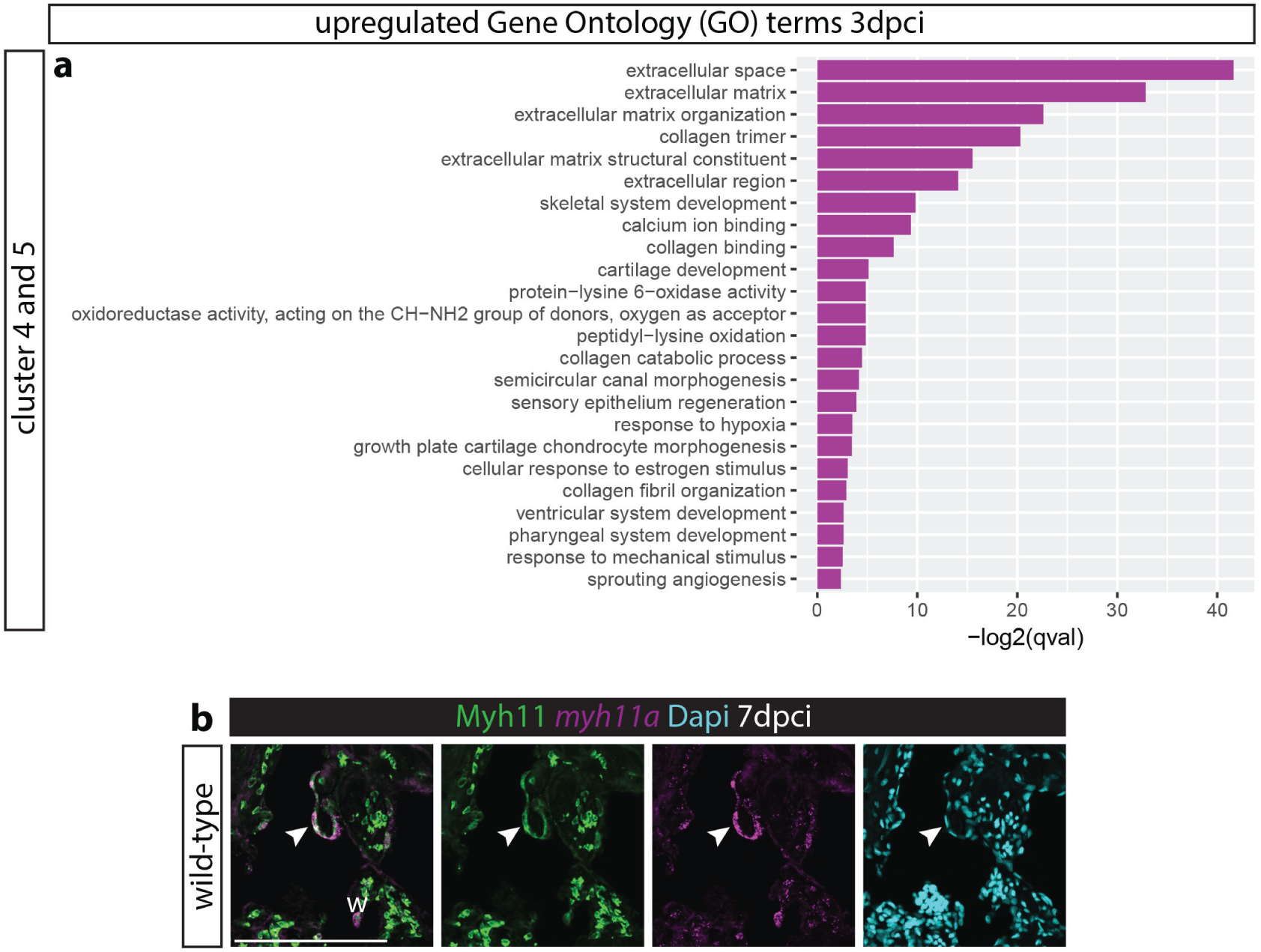
Upregulation of extracellular matrix genes in the runx1-citrine positive endocardium after injury. a, GO term analysis shows strong upregulation of GO terms associated with extracellular matrix formation in cluster 4 and 5 of the *citrine/runx1;mcherry/kdrl* positive cells after injury. b, immunohistochemistry for Myh11 combined with *in situ* hybridisation for *myh11a* and nuclear marker Dapi. Arrowhead points to overlap of myh11a mRNA with Myh11 protein, indicating specific binding of the Myh11 antibody. w, wound. Scale bars depict 100 μm.

